# Structural basis for recognition of unfolded proteins by the ER stress sensor ERN1/IRE1α

**DOI:** 10.1101/2023.06.20.545791

**Authors:** Mariska S. Simpson, Heidi De Luca, Sarah Cauthorn, Phi Luong, Namrata D. Udeshi, Tanya Svinkina, Stefanie S. Schmeider, Steven A. Carr, Michael J. Grey, Wayne I. Lencer

**Author notes:** These authors contributed equally to this work (Michael J. Grey, Wayne I. Lencer). Division of Gastroenterology, Beth Israel Deaconess Medical Center; Boston MA, 02115, United States of America;.

## Abstract

IRE1α is an endoplasmic reticulum sensor that recognizes misfolded proteins to activate the unfolded protein response (UPR). We used cholera toxin (CTx), which activates IRE1α in cells, to understand how unfolded proteins are recognized. In vitro, the A1 subunit of CTx (CTxA1) bound IRE1α lumenal domain (IRE1α_LD_). Global unfolding was not required. Instead, IRE1α_LD_ recognized a 7-residue motif within a metastable region of CTxA1 that was also found in microbial and host proteins involved in IRE1α activation. Binding mapped to a pocket on IRE1α_LD_ normally occupied by a segment of the IRE1α C-terminal flexible loop implicated in IRE1α regulation. Mutation of the recognition motif blocked CTx-induced IRE1α activation in live cells. These findings describe a mechanism for substrate recognition by IRE1α that induces the UPR.

## Introduction

IRE1α (encoded by ERN1) is one of the three proteins that monitor protein folding in the endoplasmic reticulum (ER) of mammalian cells to regulate a key cell-protective stress response termed the "Unfolded Protein Response" (UPR) (*1, 2*). The UPR adjusts the functional capacity of the ER by modulating protein translation, expansion of the ER, chaperone-dependent protein folding, and ER associated degradation (ERAD) to remove misfolded proteins (*3, 4*). Upon sensing misfolded proteins, IRE1α forms dimers or small oligomers (*5, 6*) that induce the UPR through its dual kinase-endonuclease cytosolic effector domain, primarily by splicing the mRNA for the transcription factor XBP1s (*7–9*). Deficiencies in the UPR contribute to human disease (*10–12*), but exactly how IRE1α recognizes misfolded proteins to induce this adaptive response is not clear.

IRE1α has been shown to bind unfolded proteins and peptides in vitro (*13, 14*) and misfolded proteins and peptides in cells (*15, 16*). One hypothesis is that misfolded proteins bind directly to a MHC class I-like groove on the IRE1α_LD_ surface to stabilize the active conformation (*14, 17–19*), though other binding sites have not been ruled out (*13, 20*). In addition, little is known about the features of misfolded proteins that enable recognition by IRE1α.

We addressed these problems using cholera toxin (CTx). CTx has evolved to enter the ER lumen of host cells where a portion of the toxin (the CTxA1-chain) completely unfolds to hijack ERAD and thread across the ER-limiting membrane into the cytosol for activation of cAMP and the fluid secretion that underlies the diarrhea of cholera (*21*). Somewhere in this process, the CTxA1 chain activates IRE1α (*22–24*). This provides a physiologic model to investigate the basis for how IRE1α senses unfolded proteins.

## Results

### CTxA1 binds directly to IRE1α_LD_

We first confirmed that CTxA1 activates IRE1α from within the ER of affected cells (*22–24*). Polarized T84 intestinal epithelial cells treated with CTx had increased IRE1α activation as assessed by XBP1s mRNA compared to cells treated with media or CTxB alone (Fig. 1A). XBP1 splicing was specific for IRE1α as it was blocked by the IRE1α endonuclease inhibitor 4μ8C (*25*). Similar results were found in other cell lines (fig. S1A,B) and mouse ileal tissue (fig. S1C). Activation of IRE1α by CTx occurred within the ER lumen as a mutant toxin with an A1 subunit that cannot retro-translocate into the cytosol (CTxA R192G) (*26*) still induced XBP1 splicing (fig. S1D). And treatment with forskolin (FSK), which mimics the toxic effect of CTx in the cytosol of host cells by maximally activating adenylate cyclase, did not induce XBP1 splicing as effectively (fig. S1E). We also tested for an interaction between IRE1α and CTxA1 by monitoring toxin-induced fluid secretion using primary colon epithelial organoid cultures. Both CTx and FSK caused organoid swelling due to cAMP-induced fluid secretion (Fig. 1B), but inhibition of IRE1α with 4μ8C affected only the swelling caused by CTx (Fig.1B). This result implicates IRE1α in the pathogenesis of CTx-induced disease.

**Fig. 1.**
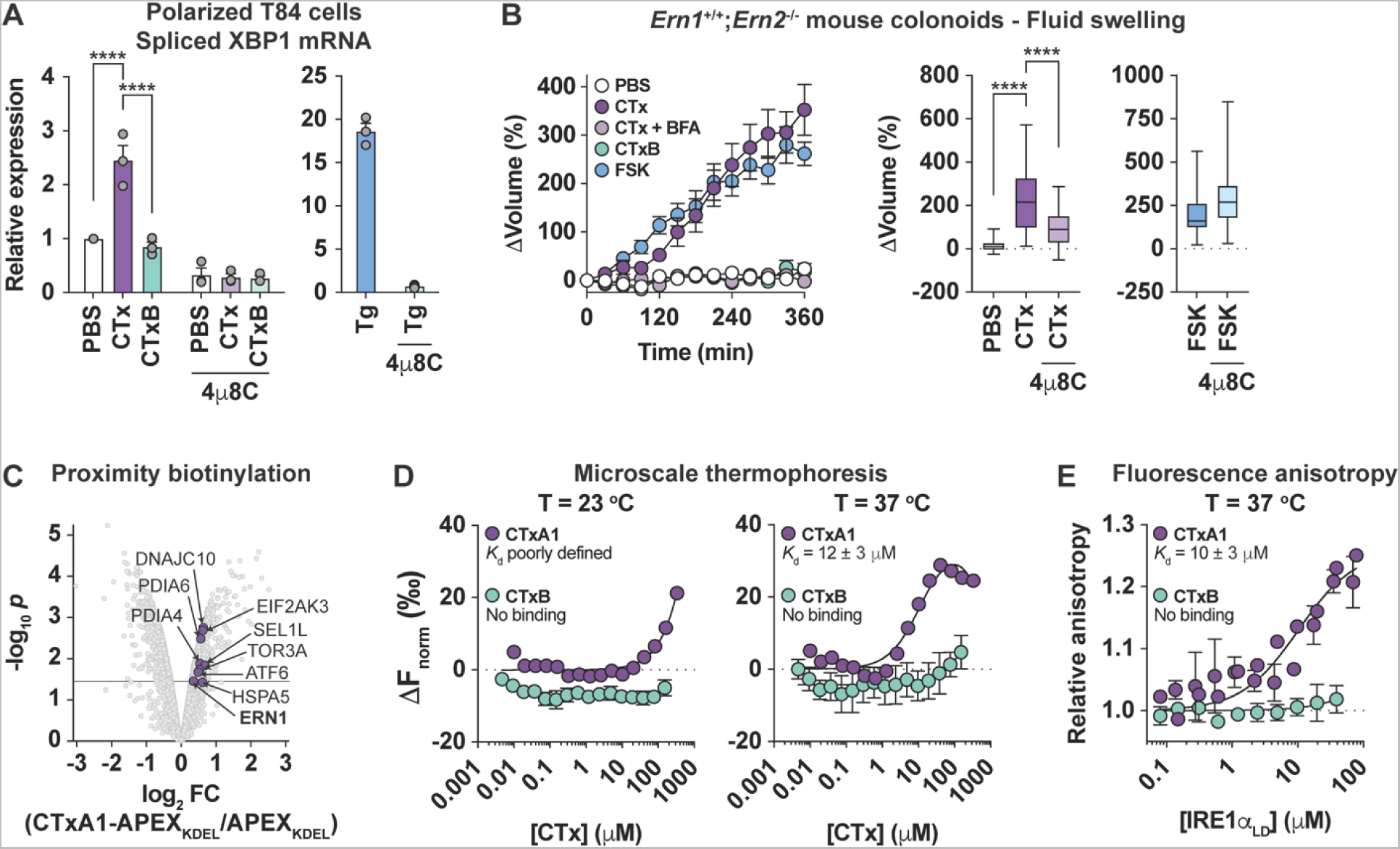
CTx A subunit activates IRE1α in the ER of live cells and binds IRE1α_LD_ in vitro. (**A**) Relative expression of spliced XBP1 mRNA assayed by qPCR from polarized T84 cell monolayers treated with PBS (white bars), CTx (purple bars), or CTxB (green bars) in the absence or presence of the IRE1α inhibitor 4μ8C. Cells treated with thapsigargin (Tg, blue bars) to induce ER stress were a positive control. Symbols represent independent experiments (n = 3), and bars represent mean ± SEM. Mean values were compared by two-way ANOVA. (**B**) cAMP-dependent fluid secretion assayed in primary mouse colonoids. (B, left panel) Time course for change in volume of colonoids treated with PBS (negative control), CTx, CTx plus brefeldin A (BfA) to block transport into the ER (*55*), CTxB, or the adenylyl cyclase agonist forskolin (FSK, positive control). Symbols represent mean ± SEM for at least 10 individual colonoids from two independent experiments. (B, right panel) Box plot represents the change in colonoid volume at t = 240 min compared to t = 0 min for at least 20 individual colonoids from two independent experiments. Mean values were compared by one way ANOVA. (**C**) Volcano plot showing enrichment of biotinylated proteins for cells expressing CTxA1-APEX_KDEL_ compared to cells expression APEX_KDEL_. ER proteins known to be involved in CTx pathophysiology and ER stress sensors are shown as purple symbols. The line represents an adjusted *p* value of 0.01. (**D**) Response curves for binding of (purple symbols) CTxA1 or (green symbols) CTxB to IRE1α_LD_ measured by microscale thermophoresis at (left panel) 23 °C or (right panel) 37 °C. Symbols represent mean ± SEM of 4 independent measures and lines represent nonlinear fit of one-site binding model. (E) Response curve for binding of IRE1α_LD_ to fluorescently labeled (purple symbols) CTxA1 or (green symbols) CTxB measured by fluorescence anisotropy at 37 °C. Symbols represent mean ± range from 2 independent experiments. Lines represent nonlinear fit of a one-site binding model.

To determine how CTx activates IRE1α, we first took an unbiased cell-based proximity labeling approach using ascorbate peroxidase (APEX2, (*27*)) fused to CTxA1 (fig. S2). After inducing proximity biotin-labelling, biotinylated proteins were isolated and identified by liquid-chromatography-tandem mass spectrometry (LC-MS/MS) using a tandem mass tag (TMT) strategy for relative quantification. In cells expressing the ER-targeted CTxA1-APEX2, IRE1α was among the ER proteins selectively enriched in biotin labeling compared to control cells expressing ER-localized APEX2 alone. Substantiating this result, other ER proteins implicated in CTx pathophysiology were biotin-labeled at similar levels of enrichment, including components of the ERAD machinery SEL1L, ERdj5/DNAJC10, and Torsin A/TOR3A (*28–30*), and the ER chaperones BiP/HSPA5, PDIA4, and PDIA6 (*26, 31*) (Fig. 1C). Thus, CTxA1 partitions within the ER lumen of live cells in proximity of IRE1α (<20 nm) suggesting that the molecules may interact.

To test for a direct interaction, we purified recombinant IRE1α_LD_, CTxA1, and CTxB and measured binding in solution by microscale thermophoresis (MST). At 23 °C, increasing concentrations of CTxA1 resulted in a change in the MST response (ΔF_norm_), but only at very high concentrations indicating a weak interaction (Fig. 1D, left panel). Since CTxA1 becomes progressively destabilized at physiologic temperatures (*32*), we also measured binding at 37 °C. Under these conditions, CTxA1 bound IRE1α_LD_ with low micromolar affinity (Fig. 1D, right panel), comparable to the apparent affinities measured for other unstable proteins that bind IRE1α_LD_ (*14, 15*). These results were validated with a fluorescence anisotropy assay, where IRE1α_LD_ bound CTxA1 with a similar affinity at 37 °C (Fig. 1E). CTxB, however, did not bind IRE1α_LD_ in either assay (Fig. 1D,E) demonstrating specificity for the CTxA1-IRE1α_LD_ interaction. Thus, IRE1α_LD_ directly and selectively binds an unstable conformation of CTxA1.

### IRE1**α** detects a metastable region of CTxA1

We then asked if global unfolding of CTxA1 was required for recognition by IRE1α_LD_. At physiologic temperatures, CTxA1 is stabilized by assembly with CTxB in the intact holotoxin (*32*). As reported previously (*33*), we found CTxA had a melting temperature of ∼50 °C in the intact holotoxin (Fig. 2A, left panel) and no loss of secondary structure when heated to at least 42 °C (fig. S3). As such, we predicted that if global unfolding was required for binding to IRE1α, stabilization of CTxA1 by assembly in the holotoxin would impair binding. We found however that CTx holotoxin still bound IRE1α_LD_ (Fig. 2A, right panel).

**Fig. 2.**
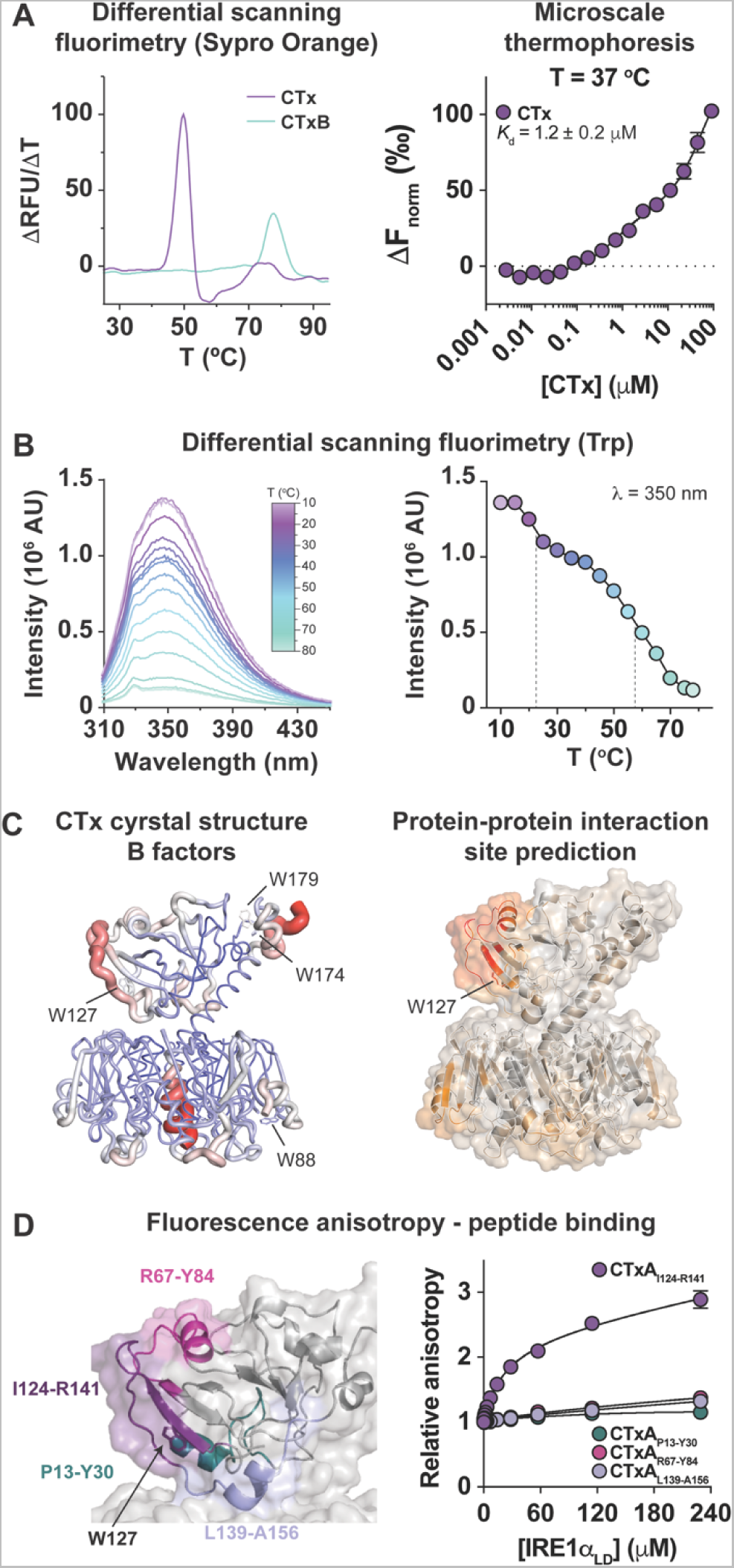
IRE1α_LD_ detects a metastable region of CTxA1 without global unfolding. (**A**, left panel) Thermal denaturation profile for CTx (purple line) and CTxB (green line) measured by differential scanning fluorimetry with Sypro Orange. Data are shown as the first derivative of the fluorescence intensity versus temperature, with peaks representing melting transitions. (A, right panel) Response curve for CTx holotoxin binding to IRE1α_LD_ measured by MST at 37 °C. Symbols represent mean ± range from 2 independent experiments. (**B**, left panel) Intrinsic tryptophan fluorescence emission scans measured for CTx as a function of temperature. (B, right panel) Thermal denaturation profile is shown for the fluorescence intensity (λ_em_ = 350 nm) as a function of temperature. Thermal transitions with approximate midpoints of 22.5 °C and 57.5 °C are indicated by dashed lines. (**C**) Ribbon diagrams of CTx crystal structure (pdb 1xtc, (*34*)) colored by (left panel) crystallographic b-factor from low (blue, thin) to high (red, thick) and (right panel) protein-protein interaction site prediction score from low (gray) to high (red). (**D**, left panel) Close-up view of CTxA1 ribbon diagram indicating CTxA1 peptides tested for binding to IRE1α_LD_. (D, right panel) Response curves for binding of IRE1α_LD_ to fluorescently labeled CTxA1 peptides measured by fluorescence anisotropy. Symbols represent mean ± SEM from 3 independent experiments and the lines represent nonlinear fit of a one site binding model to the experimental data.

To identify regions of CTxA1 that may be recognized by IRE1α in the intact toxin, we monitored unfolding by intrinsic tryptophan fluorescence. CTxA1 has three Trp residues (W127, W174, and W179) that can report on local conformation with loss of fluorescence intensity upon unfolding with increasing temperature (Fig. 2B, left panel). Two prominent thermal transitions in the fluorescence intensity (λ_em_ = 350 nm) were observed: a high temperature transition around 55 °C consistent with global unfolding of CTxA and a low temperature transition around 22.5 °C indicative of localized conformational change (Fig. 2B, right panel).

While all three Trp sidechains in CTxA1 are partially or fully buried in the CTx structure (pdb 1xtc, (*34*)), W127 resides in an edge strand of a β-sheet adjacent to a flexible loop that extends along the edge of the A1 subunit (Fig. 2C, left panel). This region (W127-R141) was computationally identified as a potential site mediating protein-protein interactions (Fig. 2C, right panel) and previously implicated in binding IRE1α (*22, 23*). To test if this segment bound IRE1α_LD_, we generated fluorescently labeled peptides of CTxA1_I124-R141_ and peptides surrounding this site (Fig. 2D, left panel). As measured by fluorescence anisotropy, IRE1α_LD_ bound CTxA1_I124-R141_ with an apparent affinity similar to CTxA1 (Fig. 2D, right panel; *K*_d_ = 44 ± 5 μM). IRE1α bound less well to peptides surrounding this site, and it did not bind a control peptide (CTxA1_P13-Y30_) representing a region buried in the CTxA1 structure. These results indicate that IRE1α_LD_ recognizes residues I124-R141 as part of a metastable region of CTxA1 that samples locally unfolded conformations at physiologic temperatures.

### CTxA1 binds a protected pocket on IRE1**α**_LD_ distal from the MHC-like groove

To determine where CTxA1 binds on IRE1α_LD_, we first tested the MHC-like binding groove thought to be the site where misfolded proteins bind IRE1α (Fig. 3A). Mutation of residues in the groove (K121A, Y161A, and Y179A) previously implicated in peptide binding for yeast Ire1p (*17*) destabilized the recombinant human IRE1α_LD_, and this undermined their use for in vitro binding studies. As an alternative, we expressed and purified the IRE1α_LD_ D123P mutant that disrupts IRE1α_LD_ dimerization and assembly of the MHC-like groove (*35*). IRE1α_LD_ D123P eluted later than WT IRE1α_LD_ on gel filtration, consistent with impaired dimerization (fig. S4). However, the D123P mutation did not impair binding to CTxA1 (Fig. 3B, left panel) or to the CTxA1_I124-R141_ peptide (Fig. 3B, right panel). Thus, CTxA1 does not bind within the MHC-like groove.

**Fig. 3.**
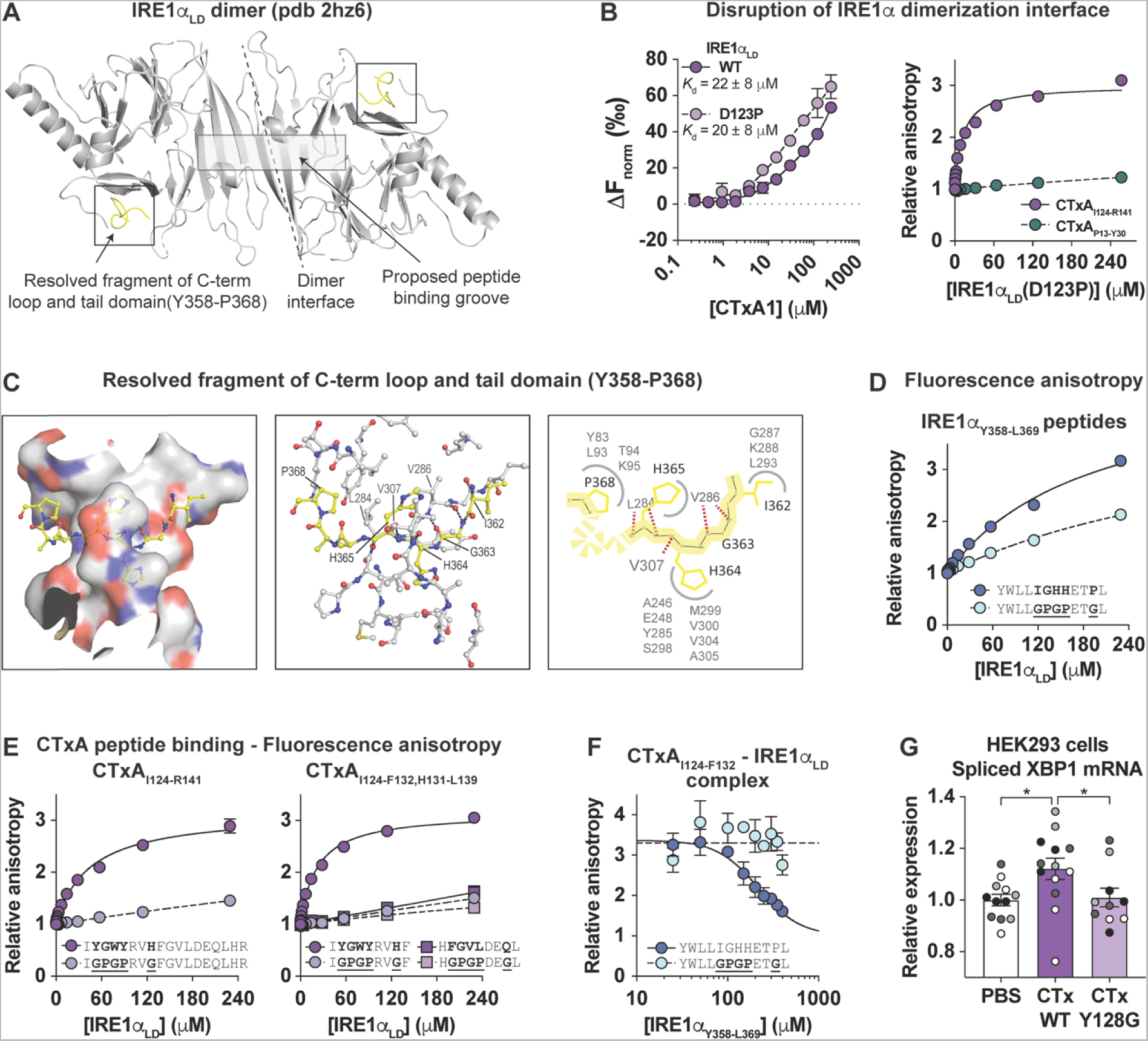
CTxA1 binds a protected pocket on the C-terminus of IRE1α_LD_ distal from the dimerization interface and MHC-like groove. (**A**) Cartoon of IRE1α_LD_ crystal structure (pdb 2hz6, (*35*)). The dimerization interface, MHC-like groove, and resolved segment (residues 358-368) of C-terminal flexible loop and tail domain are indicated. (**B**, left panel) Response curves for CTxA1 binding to WT (dark purple symbols) and D123P (light purple symbols) IRE1α_LD_ measured by MST at 37 °C. Symbols represent mean ± range for 2 independent measures, and lines represent nonlinear fit of a one-site binding model. (B, right panel) Response curves for binding of IRE1α_LD_(D123P) to indicated CTxA1 peptides measured by fluorescence anisotropy. Symbols represent measures from a single experiment, and lines represent nonlinear fit of a one-site binding model. (**C**, left and middle panels) Close-up structural representations of the interactions between IRE1α residues Y358-P368 with a pocket on the lumenal domain. Residues Y358-P368 are shown in ball-and-stick representation with carbon, nitrogen, and oxygen atoms colored yellow, blue, and red, respectively. Backbone hydrogen bonds are shown as black dashed lines. (C, right panel) Schematic illustrating the hydrogen bonding and side chain interactions. (**D, E**) Response curves for binding of IRE1α_LD_ to fluorescently labeled (**D**) IRE1α_Y358-L369_ peptides and (**E**, left panel) CTxA_I124-R141_, (right panel, circles) CTxA_I124-F132_ and (right panel, squares) CTxA_H131-L139_ peptides measured by fluorescence anisotropy; darker and lighter shade symbols represent binding to peptides with wild type and mutated recognition motifs, respectively. Symbols represent mean ± SEM for 3 independent measures, and lines represent nonlinear fit of a one-site binding model. (**F**) Response curve for displacement of IRE1α_LD_ from fluorescently labeled CTxA_I124-F132_ by competition with WT (dark blue symbols) or mutant (light blue symbols) IRE1α_Y358-L369_ peptides. Symbols represent mean ± SEM for 3 independent experiments and lines represent best fit to either a inhibitor versus response model (solid line) or a horizontal line (dashed line). (**G**) Bar graph shows relative expression of spliced XBP1 mRNA in HEK293 cells treated with PBS, WT CTx, or CTx with Y128G mutation in A subunit. Different color symbols represent 4 independent experiments each with 2-3 replicate samples within an experiment. Bars represent mean ± SEM, and mean values were compared by one way ANOVA.

To identify where CTxA1 might bind, we inspected the IRE1α_LD_ crystal structure. We noted a segment of the C-terminal flexible loop and tail region (residues 358-YWLLIGHHETP-368), resolved in two different crystal structures, docked within a pocket on the lumenal domain distal from the primary dimerization interface and MHC-like groove (Fig. 3A, pdb 2hz6 and 6shc) (*13, 35, 36*). Residues Y358-P368 adopted an extended sheet-like conformation anchored by backbone hydrogen bonds (G363-NH:V286-O, G363-O:V286-NH, H364-O:V307-NH, H365-NH:L284-O, and H365-O:L284-NH), with the side chain of H364 extending into a deep cavity and the side chains of I362 and P368 occupying surface indentations (Fig. 3C). When tested in vitro, IRE1α_LD_ bound a synthetic peptide of this segment (IRE1α_Y358-L369_, Fig. 3D dark blue symbols), and binding was impaired by targeted mutations that disrupted the backbone hydrogen bonding (G363P/H365P) and side chain interactions (I362G/H364G/P368G, Fig. 3D light blue symbols).

Notably, the CTxA1_I124-R141_ peptide that bound IRE1α_LD_ (124-I**<u>YGWY</u>**RV**<u>H</u>**FGVLDEQLHR-141) has features similar to the endogenous IRE1α_Y358-P368_ segment (358-YWLL**<u>IGHH</u>**ET**<u>P</u>**-368) and could mimic the backbone and side chain interactions required for recognition by IRE1α_LD_ (Table 1). Consistent with this idea, we found that introduction of the GPGPxxG mutations in CTxA1’_I124-R141_ peptide abolished binding (Fig. 3E, left panel). In addition, binding was detected only for the N-terminal portion of the peptide which contained the recognition motif (CTxA1_I124-F132_) but not for the C-terminal half (CTxA1_F132-R141_, Fig. 3E, right panel), thus defining YGWYRVH as the CTxA1 sequence recognized by IRE1α.

**Table 1.** Analysis of IRE1α recognition motif.

| **Motif position** | **P1** | **P2** | **P3** | **P4** | **P5** | **P6** | **P7** |
| --- | --- | --- | --- | --- | --- | --- | --- |
| IRE1α<sub>I362-P368</sub> | Ile | Gly | His | His | Glu | Thr | Pro |
| Backbone | - | ++ | + | ++ | - | - | - |
| Side chain | + | - | +++ | + | - | - | + |
| Requirements | bulky<br>ring<br>aliphatic<br>polar<br>Not Gly | small<br>Not Pro | bulky<br>ring<br>polar<br>Not Gly | bulky<br>ring<br>polar<br>Not Gly<br>or Pro | any | any | bulky<br>ring<br>aliphatic<br>polar<br>Not Gly |
| CTxA<sub>Y125-H131</sub> | Tyr | Gly | Trp | Tyr | Arg | Val | His |
| LTxA<sub>Y124-N131</sub> | Tyr | Gly | Trp | Tyr | Arg | Val | Gln |
| ORF8<sub>Y42-R48</sub> | Tyr | Ser | Lys | Trp | Tyr | Ile | Arg |
| MOG<sub>V66-P72</sub> | Val | Gly | Trp | Tyr | Arg | Ser | Pro |
| IgG C<sub>H1</sub><sub>V39-A45</sub> | Val | Ser | Trp | Gln | Ser | Gly | Ala |
| IgG C<sub>H2</sub><sub>F158-G164</sub> | Phe | Gln | Trp | Tyr | Val | Glu | Gly |
| IgG C<sub>H3</sub><sub>V262-G268</sub> | Val | Glu | Trp | Glu | Ser | Gln | Gly |
| IgG V<sub>LL33-K39</sub> | Leu | Ala | Trp | Tyr | Gln | Gln | Lys |

We next performed a competition assay to confirm that CTxA1_I124-F132_ bound to the same site on IRE1α_LD_ as the endogenous IRE1α_Y358-P368_ segment. Binding of IRE1α_LD_ to fluorescently labeled CTxA1_I124-F132_ was blocked by the IRE1α_Y358-L359_ peptide (Fig. 3F, dark blue symbols) but not by the mutated peptide with impaired binding (Fig. 3F, light blue symbols). The apparent inhibitory constant (220 ± 30 μM) was similar to the binding affinity for IRE1α_Y358-L369_ (190 ± 20 μM). Thus, CTxA1 binds to the same pocket located distally from the MHC-like groove that is normally occupied by the endogenous IRE1α_Y358-P368_ fragment.

To link such binding with IRE1α activation in live cells, we expressed and purified a mutant CTx with a single Y128G substitution in the binding motif (YGW**<u>G</u>**RVH, CTxA Y128G). A peptide with this point mutation had impaired binding to IRE1α_LD_ in vitro (fig. S5). And HEK293 cells treated with the mutant toxin had impaired activation of IRE1α as assessed by XBP1 splicing (Fig. 3G). Thus, binding of CTxA1 to the IRE1α_LD_ pocket normally occupied by the IRE1α_Y358-P368_ fragment activates IRE1α in intact cells.

### Other proteins implicated in IRE1**α** activation contain a recognition motif similar to CTxA1

To test if the features of the **<u>IGHH</u>**ET**<u>P</u>** and **<u>YGWY</u>**RV**<u>H</u>** sequences that bind IRE1α reflect a general rule for a recognition motif, we searched for similar sequences in other proteins that activate IRE1α signaling. We found similar sequences in: the Heat-labile enterotoxin from *Escherichia coli*, which is structurally and functionally related to CTx and also found to activate IRE1α in intestinal epithelial cells (*37*) (LTx: 124-I**<u>YGWY</u>**RV**<u>N</u>**F-132); the ORF8 accessory protein of SARS-CoV2, which accumulates in the ER of infected cells and activates IRE1α (*38, 39*) (ORF8: 39-IHF**<u>YSKW</u>**YI**<u>R</u>**-48); and the myelin oligodendrocyte glycoprotein, which is involved in myelin sheath maintenance and implicated in IRE1α activation (*40*) and the pathophysiology of multiple sclerosis (MOG: 64-ME**<u>VGWY</u>**RS**<u>P</u>**FS-74). In all cases, peptides corresponding to these sequences from LTx, ORF8, and MOG bound IRE1α_LD_ in vitro, and introduction of the GPGPxxG mutation disrupted binding (Fig. 4A-C).

**Fig. 4.**
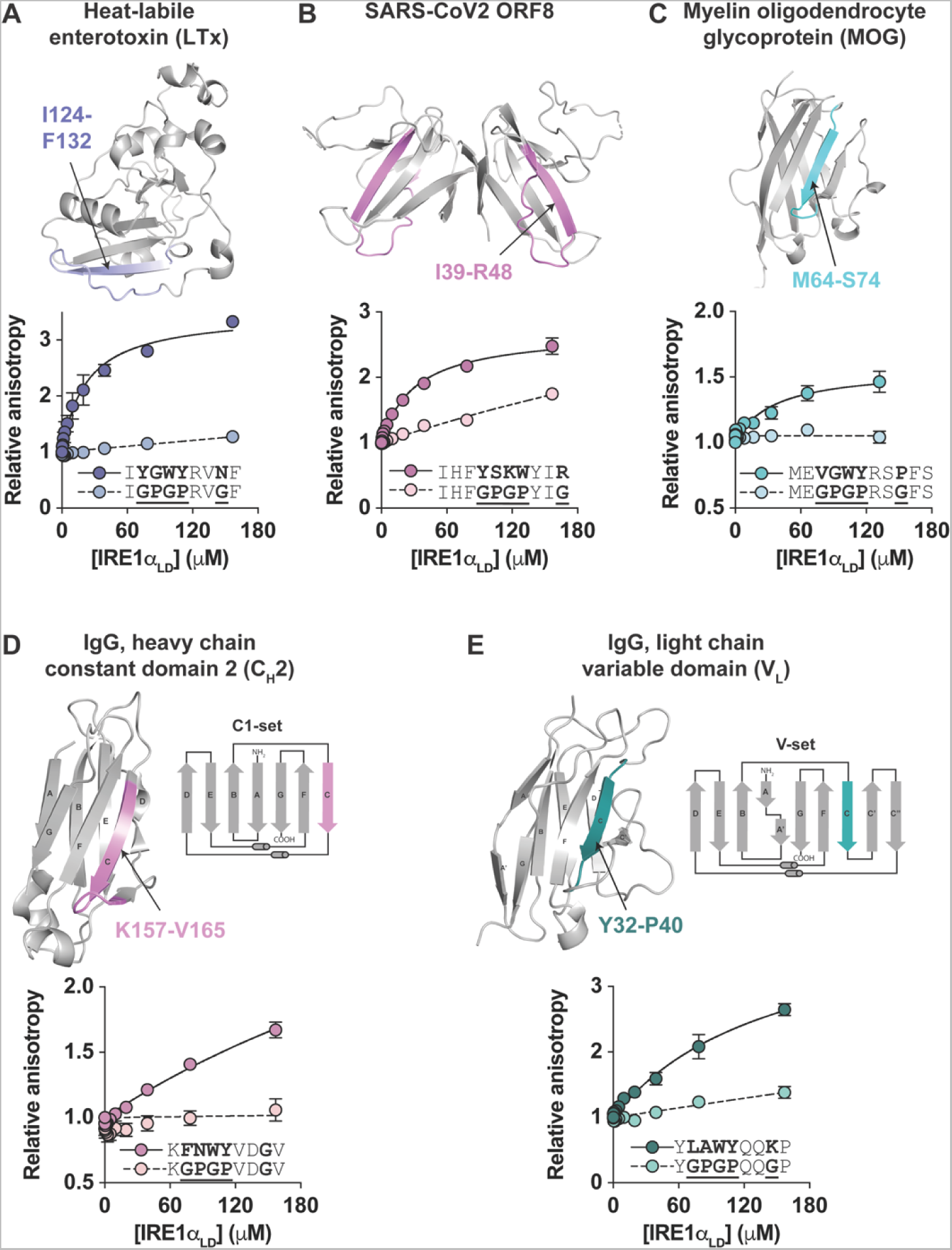
Recognition motif in other proteins that activate IRE1α. Cartoon representation of crystal structure and response curve for IRE1α_LD_ binding to fluorescently labeled peptides derived from (**A**) heat-labile enterotoxin (LTx), (**B**) SARS-CoV2 ORF8, (**C**) human myelin oligodendrocyte glycoprotein (MOG), (**D**) human IgG heavy chain constant domain 2 (C_H_2), and (**E**) human IgG light chain variable domain (V_L_) as measured by fluorescence anisotropy. The peptides sequences tested are indicated and shaded on the crystal structure. Symbols represent mean ± SEM for 3 independent measures, and lines represent nonlinear fits of one-site binding models to experimental data. A schematic illustration of β-sheet topology is shown for C_H_2 and V_L_ domains.

Each of the binding sequences that we identified were found in the edge strands of β-sheet secondary structure, similar to CTxA1. We noted that ORF8 and MOG sequences were in strands within β-sheets of immunoglobulin (Ig)-like folds. This led us to ask if antibody Ig domains contain a similar recognition motif. We found that three Ig domains comprising the constant region of human IgG heavy chain (C_H_1, C_H_2, and C_H_3) have a putative IRE1α recognition motif located in the edge strand (strand C) of one β-sheet of the C1-set Ig fold (Fig. 4D and fig. S6). And an Ig domain from the variable region of the light chain also has a putative motif located in strand C. When tested in vitro, the relevant peptides from C_H_1, C_H_2, C_H_3, and V_L_ all bound IRE1α_LD_ to varying extents (Fig. 4D,E and fig. S6), and GPGPxxG mutations in the C_H_2 and V_L_ peptides impaired their binding as predicted by our results (Fig. 4D,E). Thus, a diverse set of microbial and mammalian proteins implicated in IRE1α activation all have an IRE1α recognition motif located in edge strands of β-sheets.

## Discussion

### Implications for sensing misfolded proteins by IRE1α

Our results support a mechanism of IRE1α activation that involves direct binding between metastable regions of substrate proteins and the IRE1α_LD_. It is notable that CTxA1 does not bind in the MHC-like groove that extends across the IRE1α_LD_ dimer interface (*17*). This is consistent with an interpretation of the human IRE1α crystal structure where the opening of the groove is constrained (*35*). Although an allosteric mechanism to overcome the structural constraint has been proposed (*14*), introduction of an intermolecular disulfide to covalently close the groove did not block binding of a peptide derived from myelin protein 0 (MPZ) (*13*)—a peptide shown previously to bind IRE1α_LD_ (*14*). We also detected binding of MPZ peptide to IRE1α_LD_ (fig. S7). The MPZ peptide contains an IRE1α recognition motif, and mutation of the motif blocked binding (fig. S7). Thus, the MPZ peptide binds to IRE1α_LD_ at the same site as CTxA distal from the MHC-like groove and normally occupied by IRE1α residues Y358-P368.

Binding of misfolded proteins to this protected pocket on IRE1α_LD_ is also supported by a structure of the closely related sensor PERK/EIF2AK3. That structure revealed an exogenous peptide (P16) bound to a similar site on PERK_LD_ (fig. S8A, (*20*)). The peptide occupying the PERK binding pocket, however, lacks the IRE1α recognition motif, and we found that it did not bind IRE1α_LD_ in vitro (fig. S8B)—implying differences in specificity for substrate recognition by the IRE1α and PERK sensors. We emphasize here that other sequences or structural motifs may bind different sites on IRE1α_LD_, including within the MHC-like groove.

Though we define a binding motif and identify a binding pocket leading to activation of IRE1α, our results do not explain how binding to this site leads to activation of the endonuclease domain. Based on observations from previous studies (*5, 14*), our results do, however, suggest that displacement of IRE1α residues Y358-P368 from the binding pocket may affect IRE1α activation by regulating dimer/oligomer assembly. Activation of IRE1α likely involves a variety of oligomeric states required for auto-phosphorylation and physiologic signaling (*5*). The decisive factors driving assembly originate in IRE1α_LD_ as it responds to lumenal factors in the ER, and the flexible loop containing the IRE1α recognition motif is implicated in regulating the oligomeric state (*14*).

Of special note is that BiP binds the IRE1α_LD_ flexible loop preceding the Y358-P368 recognition motif. In an alternative hypothesis for mechanism of IRE1α activation, the ER chaperone BiP/HSPA5 binds the loop to restrain IRE1α in an inactive state (*4, 6, 13, 41-43*). When misfolded proteins accumulate, BiP dissociates from IRE1α to chaperone client proteins; and this has been associated with IRE1α dimerization and activation (*6*). Thus, it is possible that substrate binding to the IRE1α pocket may displace the flexible loop and enable the release of BiP as a necessary step in dimer/oligomer assembly. Or alternatively, in response to unfolded protein load, dissociation of BiP from the loop may enable misfolded proteins containing the IRE1α recognition motif to displace residues Y358-P368 from the binding pocket, thus freeing the loop and tail domain to mediate assembly of the active configuration. Such mechanisms would integrate the two indirect and direct models for regulation of IRE1α signal transduction.

In all cases, the sequences we found to bind IRE1α_LD_ were derived from edge strands of β-sheets, suggesting that IRE1α may serve as a sensor of unpaired β-strands containing the appropriate recognition motif. In the case of CTx, we identified this motif in the edge strand of a metastable β-sheet fold. We hypothesize that the toxin may have evolved, at least in part, for this edge strand to dissociate from the rest of the β-sheet for recognition by IRE1α and induction of the UPR, thus enhancing CTx retro-translocation to the cytosol by amplifying ERAD. This model would apply generally to other enterotoxins and viruses co-opting the ER to invade the host cell.

In the case of endogenous proteins synthesized in the ER, we found the recognition motif in the edge strand of certain Ig domains, including those of antibodies. Uniformly, the motif located to regions of Ig domains that are dynamic and relatively unstable (*44, 45*). And given the sheet topology of Ig domains (Fig. 4D,E), strand C containing the IRE1α recognition motif must emerge in the ER lumen initially unpaired - pending synthesis of the neighboring strands, which are distant in sequence. Thus, IRE1α may recognize these strands to prevent their aggregation during translation, or to signal upregulation of protein folding capacity, or both.

The structural basis for IRE1α signaling described here may have important functional and evolutionary implications for immune cell types, especially antibody-producing plasma cells. IRE1α is highly expressed in plasma cells (*46, 47*), and IRE1α-XBP1 signaling is required for plasma cell maturation (*48–50*). We also note the strong selective pressure on IRE1α sequences across vertebrate evolution (*51*) that occurred coincident with the onset of an adaptive immune system (*52*) and the appearance of immune molecules containing Ig domains, including antibodies - the major secretory product of plasma cells. As such, our results suggest that IRE1α and antibody Ig domains may have co-evolved together - driven by the need for plasma and other immune cell types to enhance their capacity for secretory protein translation and with the metastable fold of the antibody Ig domains providing the physiological cue for IRE1α to activate the UPR. In this case, IRE1α activation would occur not only as a response to an abundance of unfolded proteins (typifying ER stress) but also as a preemptive mechanism to adapt the protein folding capacity and function of the ER in response to physiologic needs (*53, 54*).

## Materials and Methods

### Cell culture experiments

All cell lines were grown at 37 °C and 5% CO_2_. MEF and HEK293 cell lines were maintained in Dulbecco’s modified Eagle’s medium (DMEM) supplemented with 10% Fetal Bovine Serum (FBS). Human intestinal epithelial T84 cells were maintained in 1:1 DMEM/F12 media supplemented with 6% newborn calf serum. T84 cells were plated on 0.33 cm^2^ Transwell inserts (3 μm pore size polyester membranes, Corning) and allowed to polarize for 7 days. A volt/Ohm meter (EVOM, World Precision instrument) was used to measure transepithelial electrical resistance to assess monolayer formation. Treatments were applied both in the apical and basolateral compartments. Cells were pre-treated with 50 μM 4μ8c or DMSO for 30min prior to treatment with wild type or mutant CTx (3-30 nM as indicated in figure legends), CTxB (3-30 nM), thapsigargin (Tg, 0.2-3 μM), or forskolin (FSK, 10 μM) for 4 hr. Cells were washed with ice-cold PBS and used for RNA extraction.

### In vivo experiments in mice

All experimental procedures involving mice were approved by Boston Children’s Hospital Institutional Animal Care and Use committee. WT C57BL/6 mice were orally gavaged with PBS (150 μL) or CTx (50 μg in 150 μL PBS). After 4 hr, mice were euthanized, and the ileum excised and flushed with ice-cold PBS. The washed ileum was opened longitudinally, cut into several small pieces, and washed several times in PBS for RNA extraction.

### XBP1-luciferase reporter assay

XBP1 splicing reporter plasmid (pCAX-HA-2xXBP1-Luc-F) (*56*) was provided by T. Iwawaki (Kanazawa Medical University, Uchinada, Japan). pRL SV40 *Renilla* luciferase reporter vector is available commercially from Promega. HEK293 cells were seeded at 0.5x10^5^ cells/well in 96-well clear bottom white plates (Corning) and transfected with inducible luciferase reporter vectors (50 ng pCAX-HA-2xXBP1-Luc-F, 1 ng pRL-SV40-*Renilla*, and pcDNA4 empty vector), with 100 ng total DNA per well using polyethyleneimine (PEI; linear 25 kDa; Polysciences) at a DNA/PEI mass ratio of 1:3. A non-transfected control was included for correction of background luminescence. At 18-24 hr post-transfection, cells were pre-treated with 4μ8c or DMSO for 30 min before treatment with 3 μM Tg, 3 nM CTx, 3 nM CTxB or media. At 6 hr post-treatment, cells were lysed using the Dual-Glo Luciferase Assay System (Promega). Luminescence was measured using a Spark 10M plate reader (Tecan).

### Expression analysis by qPCR

RNA was extracted from cells and tissue using RNeasy Mini Kit (Qiagen) with on-column DNA digest (Qiagen) according to manufacturer’s protocol. Total RNA concentration and quality was assessed by absorbance at 260 nm and the 260/280 nm ratio, respectively. Isolated RNA (500 ng) was reverse transcribed with iScript cDNA Synthesis Kit (BioRad). qPCR was performed using primers (Table S1) and Sso Advanced Universal SYBR Green Supermix (BioRad). Samples were processed as technical triplicates with the number of independent experimental replicates reported in the figure legends. Threshold cycle (C_q_) values were measured using the CFX384 Real-Time System (BioRad). The mean expression ratio of the test sample compared to the control sample was calculated by applying the 2^-ΔΔCt^ method. The Cq values for targets were analyzed relative to the geometric mean of the *HPRT1, PPIA,* and *GAPDH* housekeeping genes.

### Mouse colonoid swelling assay

All housing and procedures involving live vertebrate animals reviewed and approved by Boston Children’s Hospital Institutional Animal Care and Use committee. Primary mouse colonoids were generated from *Ern1*^+/+^;*Ern2*^-/-^ mice (C57BL/6 background) as previously described (*57, 58*). *Ern2*-deficient mice were used as the epithelial specific paralogue IRE1β (ERN2) inhibits IRE1α endonuclease activity (*57*). Briefly, mice were euthanized, the colon excised, lumenal contents were gently removed, and the tissue flushed with ice-cold PBS. The tissue was cut open longitudinally and then into several (2-3 mm) pieces and washed several times in ice-cold PBS. Washed tissue was incubated in PBS with 10 mM EDTA for 45 min with end-over-end rotation. Epithelial cells were dissociated by vigorous shaking for 5-7 min, the supernatant containing the dissociated crypts were collected and diluted twofold with base media (Advanced DMEM/F12 supplemented with 20% FBS, 10 mM Hepes, 1X Glutamax, 1X penicillin/streptomycin). Collected cells were passed through a 100 μM strainer and filtered through a 40 μM strainer. Intact colon crypts retained on the 40 μM filter were washed and resuspended in base media and pelleted at 300 × *g* for 3 min. Crypts were resuspended in Matrigel (Corning) on ice and plated in 30 μL drops in 24-well plates. Colonoids were maintained at 37 °C and 5% CO_2_ in complete media (base media supplemented with 50% WRN-conditioned media prepared from L cells expressing Wet/R-spondin/Noggin (*59*) and 10 μM Y-27632). Media was replaced every other day and cultures were passaged by dissolving the Matrigel using Cell Recovery Solution (Corning), mechanically disrupting by pipetting, pelleting, and resuspending cells in 1.5 to 2-fold more Matrigel.

For experiments, *Ern1*^+/+^;*Ern2*^-/-^ colonoids were pre-treated with 50 μM 4μ8c, 5 μg/mL BfA, or DMSO for 30 min prior to treatment with 10 nM CTx, 10 nM CTxB, 100 μM FSK, or media. Organoid swelling was monitored using a BioTek Cytation 5 automated microscopy plate reader set at 37 °C and 5% CO_2_. Brightfield images (2.5x objective) were collected in a 5x4 grid over each Matrigel drop every 15 min. Images were processed with ImageJ. Colonoids with a starting volume less than 2 x 10^4^ pixels^3^ were analyzed for swelling by measuring the change in diameter of the largest cross-section at each time point.

### Ascorbate peroxidase (APEX2) proximity labelling assay

#### Expression of APEX constructs in HEK293T cells

Ascorbate peroxidase (APEX2) was fused to the C-terminus of the CTxA1 chain containing a signal sequence for expression in the ER and a KDEL motif for retention in the ER. As a negative control, we used an ER-targeted APEX2 construct with a C-terminal KDEL motif. Constructs were expressed by transient transfection in HEK293T cells. Localization of both constructs to the ER was verified by confocal microscopy using ER-mCherry to label the ER (fig. S2A). For experiments, constructs were expressed for 24 hr, incubated with phenol-biotin for 30 min, and then exposed to H_2_O_2_ for 1 min to activate APEX2 and induce proximity biotin-labeling. Lysis and streptavidin pull-down steps were performed, as previously described (*27*). An additional desalting step was performed to remove excess biotin from the lysate before addition to streptavidin magnetic beads.

#### On-bead trypsin digestion of biotinylated proteins

Samples collected and enriched with streptavidin magnetic beads were washed with 200 μL of 50 mM Tris-HCl buffer (pH 7.5), and 2x with 200 μL of 50 mM Tris (pH 7.5) buffer. Samples were incubated in 0.4 μg trypsin in 80 μL of 2 M urea/50 mM Tris buffer with 1 mM DTT, for 1 hr at room temperature while shaking at 1000 rpm. Following pre-digestion, 80 μL of each supernatant was transferred into new tubes. Beads were then incubated in 80 μL of the same digestion buffer for 30 min while shaking at 1000 rpm. Supernatant was transferred to the tube containing the previous elution. Beads were washed twice with 60 μL of 2 M urea/50 mM Tris buffer, and these washes were combined with the supernatant. The eluates were spun down at 5000 × g for 30 s and the supernatant was transferred to a new tube. Samples were reduced with 4 mM DTT for 30 min at room temperature, with shaking. Following reduction, samples were alkylated with 10 mM iodoacetamide for 45 min in the dark at room temperature. An additional 0.5 μg of trypsin was added and samples were digested overnight at room temperature while shaking at 700 × g. Following overnight digestion, samples were acidified (pH < 3) with neat formic acid (FA), to a final concentration of 1% FA.

#### TMT labeling and fractionation

Desalted peptides were labeled with TMT10 reagents (ThermoFisher Scientific). Peptides were resuspended in 80 μL of 50 mM HEPES and labeled with 20 μL 25 mg/mL TMTpro18 reagents in ACN. Samples were incubated at RT for 1 hr with shaking at 1000 rpm. TMT reaction was quenched with 4 μL of 5% hydroxylamine at room temperature for 15 min with shaking. TMT-labeled samples were combined, dried to completion, reconstituted in 100 μL of 0.1% FA, and desalted on StageTips. TMT labeled peptide sample was fractionated by SCX StageTips to create 3 final fractions as previously described (*60*). Eluted peptides were dried to completion.

#### Liquid chromatography and mass spectrometry

All peptide samples were separated with an online nanoflow Proxeon EASY-nLC system (Thermo Fisher Scientific) and analyzed on an Orbitrap Lumos mass spectrometer (Thermo Fisher Scientific). Each sample was injected onto an inhouse packed 20 cm x 75 μm internal diameter C18 silica picofrit capillary column (1.9 mm ReproSil-Pur C18-AQ beads, Dr. Maisch GmbH, r119.aq; PicoFrit 10 μm tip opening, New Objective, PF360-75-10-N-5). Mobile phase flow rate was 200 nL/min, comprised of 3% acetonitrile/0.1% formic acid (Solvent A) and 90% acetonitrile/0.1% formic acid (Solvent B).

The 110-min LC–MS/MS method used the following gradient profile: (min:%B) 0:2, 1:6; 85:30; 94:60; 95:90; 100:90; 101:50; 110:50 (the last two steps at 500 nL/min flow rate). Data acquisition was done in the data-dependent mode acquiring HCD MS/MS scans (r = 50,000) after each MS1 scan (r = 60,000) using a top speed approach (cycle time 2 s) to trigger MS/MS. The maximum ion time utilized for MS/MS scans was 105 ms; the HCD-normalized collision energy was set to 38; the dynamic exclusion time was set to 45 s, Charge exclusion was enabled for charge states that were unassigned, 1 and >6.

#### Analysis of mass spectrometry data (peptide level, protein level)

Mass spectrometry data was processed using Spectrum Mill v 7.08 (<u>proteomics.broadinstitute.org</u>). For all samples, extraction of raw files retained spectra within a precursor mass range of 750-6000 Da and a minimum MS1 signal-to-noise ratio of 25. MS1 spectra within a retention time range of +/- 60 s, or within a precursor m/z tolerance of +/- 1.4 m/z were merged. MS/MS searching was performed against a human Uniprot database containing APEX2 construct sequences and common laboratory contaminants. Digestion parameters were set to “trypsin allow P” with an allowance of 4 missed cleavages. The MS/MS search included fixed modification of carbamidomethylation on cysteine. TMT10 was searched using the ‘TMT10-Full-Lys’ option. Variable modifications were acetylation and oxidation of methionine. Restrictions for matching included a minimum matched peak intensity of 30% and a precursor and product mass tolerance of +/- 20 ppm.

Peptide spectrum matches (PSMs) were validated using a maximum false discovery rate (FDR) threshold of 1.2% for precursor charges 2 through 6 within each LC-MS/MS run. Protein polishing autovalidation was further applied filter the PSMs using a target protein–level FDR threshold of zero. TMT10 reporter ion intensities were corrected for isotopic impurities in the Spectrum Mill protein/peptide summary module using the afRICA correction method which implements determinant calculations according to Cramer’s Rule. We required fully quantified unique human peptides for protein quantification. We used the Proteomics Toolset for Integrative Data Analysis (Protigy, v1.0.4, Broad Institute, https://github.com/broadinstitute/prodigy) to calculate moderated *t*-test *P* values for regulated proteins.

### Expression and purification of recombinant proteins

All plasmids used for expression of recombinant IRE1α_LD_, CTx, CTxA1, and CTxB are described in Data S1. All plasmids were constructed using standard restriction enzyme-based molecular cloning. In most cases, insert DNA was synthesized as gene blocks with 5’ and 3’ restriction sites added (IDT). Inserts and plasmids were digested with restriction enzymes (New England BioLabs), gel purified (GeneJet Gel Extraction Kit, ThermoFisher), and ligated (Quick Ligation Kit, New England BioLabs). Ligated products were transformed into NEB5alpha cells and grown on LB Agar plates with appropriate antibiotic selection. Positive clones were identified by restriction digest and sequencing. Mutations were introduced by either including the mutation in the synthesized DNA insert or by site directed mutagenesis (Agilent).

GST-IRE1α_LD_ (S24-D443) was expressed from pGEX-TEV vector in BL21(DE3) pLysS *E.coli* (Invitrogen). Expression was induced in log phase cultures (OD600 = 0.6-1.0) with 0.4 mM IPTG and incubation at 16 °C for 16-20 hr. Cultures were harvested by centrifugation at 5,000 × *g* for 10 min at 4 °C. The pellet was resuspended in 1X PBS, 0.5% Triton X-100, 1x Complete protease inhibitor (Sigma) and homogenized using Emulsiflex C5 homogenizer (Avestin) at 5,000 psi. Lysate was cleared by centrifugation at 15,000 × *g* for 10 min and supernatant collected. GST-IRE1α_LD_ was bound to glutathione agarose (GoldBio) for 1 hr at 4 °C. The slurry was transferred to Econo-Pac chromatography column (BioRad) and washed 3 times with 20 mL wash buffer (1X PBS). IRE1α_LD_ was eluted using TEV protease diluted in 2x TEV buffer (50 mM Tris pH 8, 1 mM EDTA, 2 mM DTT) overnight at 4 °C. In some cases, the sample was further purified by anion exchange chromatography. Samples were dialyzed into 50 mM Tris pH 8.0, loaded onto a HiTrap Q HP column (Cytiva) equilibrated with buffer A (50 mM Tris pH 8.0, 5 mM β-mercaptoethanol), and eluted with buffer B (Buffer A with 1 M NaCl). Fractions containing IRE1α_LD_ were identified by reducing SDS-PAGE and Coomassie staining (Fisher), pooled, concentrated (Amicon Ultra 4 mL and 0.5 mL 10 kDa, Millipore), and injected on a Superdex 200 10/300 (Cytiva) gel filtration column equilibrated with running buffer (1X PBS pH 7.4, 1 mM DTT). Fractions containing protein were identified by reducing SDS-PAGE, and bands were excised for protein identification by mass spectrometry (Taplin Mass Spectrometry Facility, Harvard Medical School).

His_6_-IRE1α_LD_ and His6-CTxA1 constructs were expressed from pET28a vector in BL21(DE3) E. coli. Expression was induced in log phase cultures with 0.4 mM IPTG and incubation at 16 °C for 16-20 hr. Bacteria cells were isolated and lysed as described above. His-tagged proteins were affinity purified by binding to cobalt resin (GoldBio) equilibrated in 1X PBS. His-tagged constructs were eluted from resin with in 50 mM Tris pH 8.0, 150 mM NaCl, and 250 mM imidazole. Samples were further purified by anion exchange chromatography and gel filtration chromatography as described above for GST-IRE1α_LD_.

CTxB was expressed from pET28a vector in Shuffle T7 Express *E. coli* (New England BioLabs). Protein expression, harvesting, and cell lysis were performed as described above. CTxB was affinity purified by binding to cobalt resin and elution with 50 mM Tris pH 8.0, 150 mM NaCl, and 250 mM imidazole. Following affinity purification, CTxB was further purified by cation exchange chromatography on a HiTrap SP HP column (Cytiva) equilibrated with buffer A (5 mM Na_2_PO_4_ pH 7.0). Bound sample was eluted with a linear gradient with buffer B (5 mM Na_2_PO_4_ pH 7.0, 1M NaCl). CTxB-containing fractions were pooled, concentrated, and further purified by gel filtration on Superdex 200 in 1X PBS pH 7.4.

Intact CTx holotoxins (WT, CTxA R192G, and CTxA Y128G) were prepared by expression of CTxA and CTxB from a bicistronic insert in pBAD expression vector. Both A and B subunits had a heat-labile enterotoxin signal peptide at the N-terminus to target expressed protein to the periplasm. Expression was induced in DH10b cells using 0.5% arabinose for 3 hr at 37 °C, or 16-20 hr at room temperature. Cultures were harvested by centrifuging at 5,000 *g* for 10 min at 4 °C. The cell pellet was resuspended in 50 mM NaHPO_4_, 300 mM NaCl, pH7.0, and 1X Complete Protease Inhibitor and treated with 0.5 mg/mL polymoxin 1 hr at room temperature to disrupt the outer membrane and release proteins from the periplasmic space. The periplasmic fraction was isolated by centrifugation at 15,000 × *g* for 10 min and supernatant collected. Intact holotoxin was affinity purification by binding to cobalt resin for 1 hr at 4°C, transferring to an Econo-Pac chromatography column, washing 3 times with 20 mL wash buffer (50 mM Na_2_HPO_4_, 300 mM NaCl, and 10 mM imidazole), and elution with 300 mM imidazole in wash buffer. Eluted protein was buffer exchanged into 50 mM Tris pH 8.5, loaded on a HiTrapQ column and eluted with a linear gradient with 50 mM Tris pH 8.5, 1 M NaCl. Pooled fractions containing both CTxA and CTxB were concentrated and loaded on a Superdex 200 gel filtration column and eluted with 1X PBS pH 7.3 to separate intact holotoxin (CTxA + CTxB) from free CTxB.

### Thermal stability assays

Thermal stability of CTx was assayed by circular dichroism (CD) spectroscopy and differential scanning fluorimetry (DSF). CD experiments were performed on a J-810 spectrometer (Jasco). Measurements were perfomred on 300 μL of 1 μM CTx in PBS. Spectra was recorded from 200 to 280 nm at temperatures of 18-65 °C; samples were equilibrated for 10 min at each temperature before measurement. Global unfolding of CTx was monitored by DSF using Sypro Orange (Thermo Fisher) on a CFX96 Real-time qPCR instrument (BioRad). Melting curves were collected in a 96-well plate with 1 μM CTx or CTxB and 5X Sypro Orange dye in PBS pH 7.4 (20 μL reaction volume). Fluorescence was measured using the FRET channel at temperatures of 25-95 °C in 0.5 °C increments and averaged over 4 independent samples.

Melting transitions were identified as the peaks of the first derivative of fluorescence intensity as a function of temperature. Localized unfolding of CTx was assayed by DSF using intrinsic tryptophan fluorescence measured on a Fluorlog-3 spectrofluorometer (Horiba) equipped with a Peltier thermostatted cuvette holder. Samples contained 1 μM CTx or CTxB in 250 μL of PBS. Tryptophan fluorescence was measured by excitation at 295 nm with emission scans from 300-500 nm at temperatures of 10 °C to 78 °C in increments of 5 °C; samples were equilibrated for 10 min at each temperature step. Fluorescence intensity at 350 nm was averaged over three independent measures and plotted as a function of temperature. Melting transitions were estimated by the first derivative of fluorescence intensity at 350 nm as a function temperature.

### In vitro binding by microscale thermophoresis (MST)

IRE1α_LD_ was labelled using the Monolith NT protein labelling Kit Red-Maleimide (NanoTemper). The concentration of the labelled IRE1α_LD_ was kept constant at 10 nM, and equal volume of the non-labelled binding partner (CTxA1, CTxB or CTx) was added as a two-fold serial dilution. Final buffer composition was 1X PBS, 1 mM DTT, 2.5% glycerol and 0.05% Tween-20. The samples were loaded into Monolith NT.115 Premium Capillaries (NanoTemper Technologies) and incubated at 23 °C or 37 °C. The MST measurement was performed using the Monolith NT.115 Pico-red (NanoTemper Technologies) at 5% LED power and medium MST power. Binding was analyzed by nonlinear fitting of a one-site (total and nonspecific) binding model to the data.

### In vitro binding by fluorescence anisotropy

His_6_-CTxA1 was labeled using Monolith NT His-Tag Labeling Kit RED-tris-NTA (NanoTemper). CTxB labeled with Alexa Fluor 647 (Invitrogen) was used as a negative control. Binding reactions were measured in 384-well black flat-bottomed assay plates in 20 μL reactions containing 50 nM of labeled protein and varying concentrations of GST-IRE1α_LD_ in 1X PBS, 1 mM DTT, 2.5% glycerol, and 0.05% Tween-20. Anisotropy was measured on a Spark 10M plate reader (Tecan) with excitation at 640 nm and emission at 685 nm and averaged over a 20 min time course. A well containing only buffer was included for background correction. Response curves were analyzed by nonlinear fitting of a one-site binding model to the experimental data.

All peptides used for binding assays (Data S1) were synthesized with a N-terminal 5-carboxyfluorescien (5-FAM) at >85% purity (GenScript); peptides used for competition studies did not have 5-FAM. Peptides were reconstituted in DMSO and further diluted in binding buffer. Binding reactions were measured in 384-well black flat-bottomed assay plates with 20 μL volume containing 50 nM of labeled peptide and varying concentrations of IRE1α_LD_ in 1X PBS, 1 mM DTT, 2.5% glycerol, and 0.05% Tween-20. Binding reactions were equilibrated at room temperature for 20 min. Fluorescence anisotropy was measured on a Spark 10M plate reader (Tecan) with excitation at 485 nm and emission at 525 nm and averaged over a 20 min kinetic time course. A well containing only buffer was included for background correction. Response curves were analyzed by nonlinear fitting of a one-site binding model to the experimental data.

Competition assays were performed by titrating varying concentrations of unlabeled IRE1α_Y358-L369_ peptides in reactions containing 50 nM labeled CTxA_I124-F132_ peptide. Equal volume of unlabeled 50 µM IRE1α_LD_ or binding buffer was added to peptides and equilibrated for 20 min. Final buffer composition was 1X PBS, 1 mM DTT, 2.5% glycerol and 0.05% Tween-20. A peptide-only control was included per datapoint that did not contain IRE1α_LD_ for background correction. Anisotropy was measured as described above.

### Prediction of CTx protein-protein interaction sites

Potential sites involved in protein-protein interactions for CTx was analyzed using the meta protein-protein interaction site predictor (meta-PPISP) server (https://pipe.rcc.fsu.edu/meta-ppisp.html, (*61*). The crystal structure of CTx holotoxin was used as the input PDB file (1xtc, (*34*)). Meta-PPISP prediction scores were mapped onto the crystal structure and amino acids were colored from low (gray) to high (orange) probability of a being a protein-protein interaction site.

### Statistical Analysis

All analyses were performed using Prism (GraphPad). The number of independent experiments is indicated in the figure legends. Figures include all independent measures shown as mean value ± SEM. Expression data is represented as fold changes (e.g., treatment compared with control) with mean values compared using unpaired t-test, one-way ANOVA, or two-way ANOVA as appropriate with correction for multiple comparisons. Significance is indicated by asterisks (* *p* < 0.05; ** *p* < 0.01; *** *p* < 0.001; **** *p* < 0.0001).

## Acknowledgments

We thank members of the Lencer Lab and Ineke Braakman (Utrecht University) for feedback throughout the course of this project; the Harvard Digestive Disease Center for services and instrumentation provided by the Epithelial Cell and Mucosal Immunology Core and the Microscopy and Histopathology Core; the Center for Macromolecular Interactions (Harvard Medical School) for access to microscale thermophoresis instrumentation; and the Taplin Mass Spectrometry Facility (Harvard Medical School) for identification/validation of purified proteins.

## Author contributions

Conceptualization: MSS, MJG, WIL

Methodology: MSS, PL, SSS, SAC, MJG, WIL

Investigation: MSS, HD, SC, PL, NDU, TS, SSS, SAC, MJG

Visualization: MSS, NDU, MJG, WIL

Funding acquisition: MJG, WIL

Project administration: MSS, MJG, WIL

Supervision: MJG, WIL

Writing – original draft: MSS, MJG, WIL

Writing – review & editing: all authors

## Data and materials availability

All data are available in the main text or the supplementary materials. All plasmids for expression of recombinant proteins will be made available upon request. The original mass spectra, peptide spectrum match results, and the protein sequence databases used for searches have been deposited in the public proteomics repository MassIVE (http://massive.ucsd.edu) and are accessible at ftp://MSV000091745@massive.ucsd.edu when providing the dataset password: proximity. If requested, also provide the username: MSV000091745. These datasets will be made public upon acceptance of the manuscript.

---

**Fig. S1.**
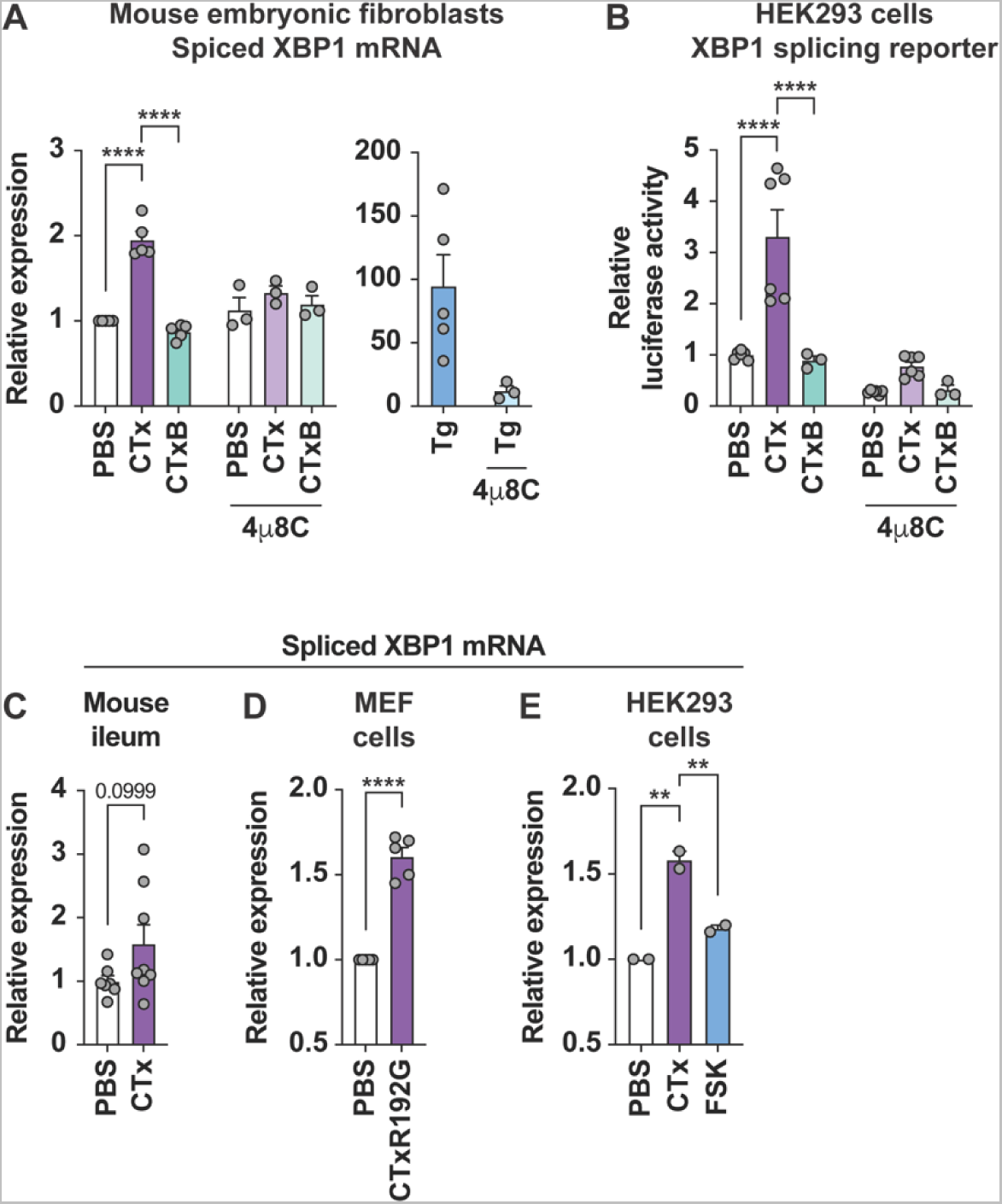
(A) Relative expression of spliced XBP1 mRNA assayed by qPCR from mouse embryonic fibroblasts (MEF) treated with PBS (white bars), CTx (purple bars), or CTxB (green bars) in the absence or presence of the IRE1α inhibitor 4μ8C. Cells treated with thapsigargin (Tg, blue bars) were included as a positive control for induction of spliced XBP1 mRNA. Symbols represent independent experiments (n = 3-5) and bars represent mean ± SEM. Mean values were compared by two-way ANOVA. (B) Luciferase activity measured in HEK293 cells transfected with XBP1 splicing luciferase reporter (*56*) and treated as in (A). Symbols represent individual wells from 2 independent experiments (3 wells per experiment) and bars represent mean ± SEM. Mean values were compared by two-way ANOVA. (C) Relative expression of spliced XBP1 mRNA assayed by qPCR from ileal tissue of mice treated with PBS (white bars) or CTx (purple bars). Symbols represent individual animals from 2 independent experiments, and bars represent mean ± SEM. Mean values were compared by unpaired t-test. (D) Same as (A) for MEFs treated with PBS or CTx contain an A subunit with R192G mutation. Symbols represent individual experiments (n = 5), and bars represent mean ± SEM. Mean values were compared by unpaired t test. (E) Relative expression of spliced XBP1 mRNA assayed by qPCR from HEK293 cells treated with PBS, CTx, or the adenylyl cyclase agonist forskolin (FSK). Symbols represent individual experiments (n = 2), and bars represent mean ± range. Mean values were compared by one-way ANOVA.

**Fig. S2.**
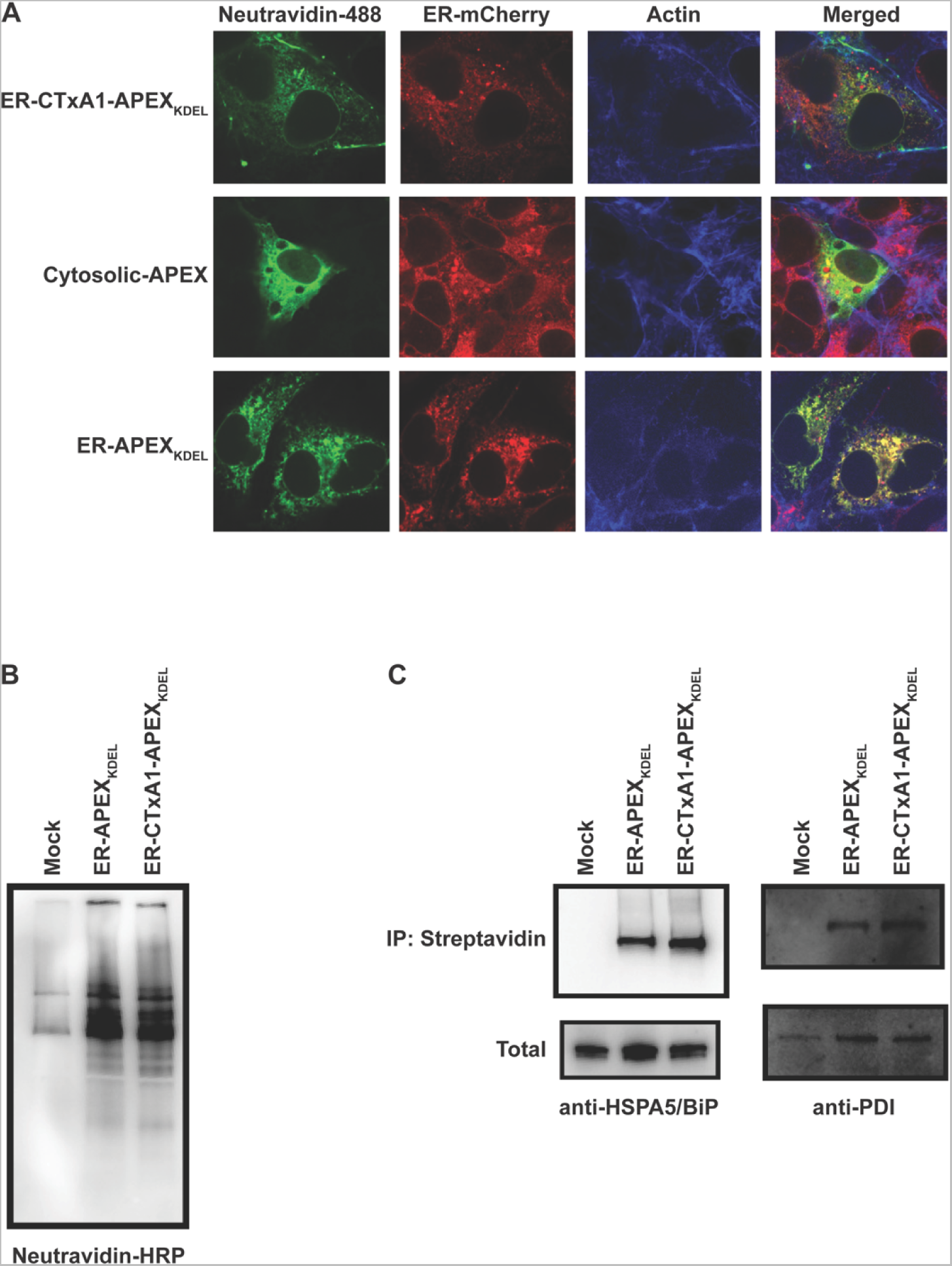
Representative micrographs of HEK293T cells expressing ER-CTxA1-APEX_KDEL_, Cytosolic-APEX, or ER-APEX_KDEL_. Cells were co-transfected with an ER-mCherry construct to label the ER. Cells were stained for biotinylated proteins and actin. Note, the co-localization of biotinylated proteins (neutravidin-488 staining) with ER-mCherry for only cells expressing ER-CTxA1-APEX_KDEL_ and ER-APEX_KDEL_ (not Cytosolic-APEX), indicating biotinylation of proteins within the ER. (B,C) Immunoblot of biotin-labeled proteins from cells expressing no APEX, ER-APEX_KDEL_, and ER-CTxA1-APEX_KDEL_. Biotinylated proteins were enriched by immunoprecipitation with streptavidin-coated beads followed by immunoblotting with (B) neutravidin-HRP, (C, left panel) anti-HSPA5/BiP, or (C, right panel) anti-PDI.

**Fig. S3.**
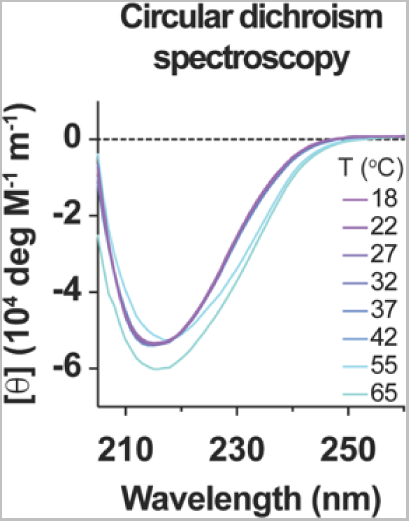
Far-UV circular dichroism spectra were measured for CTx at the indicated temperatures from 18 - 65 °C.

**Fig. S4.**
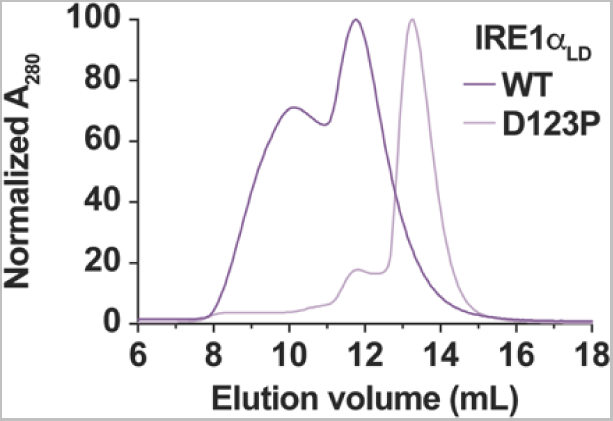
Gel filtration profiles for WT (dark purple) and D123P (light purple) IRE1α_LD_ were measured by the absorbance at 280 nm (A280). Chromatograms are shown as the normalized absorbance relative to the maximum within the trace.

**Fig. S5.**
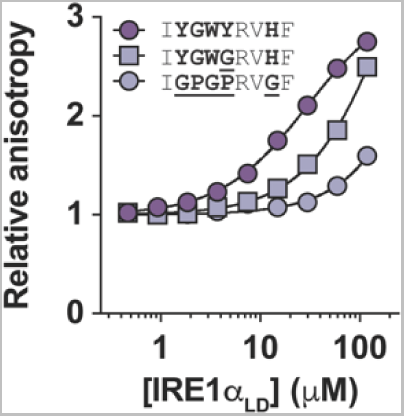
Response curve for binding of IRE1α_LD_ to CTxAI_124-F132_ peptide with WT sequence (dark purple circles), GPGPxxG mutation (light purple circles), and Y128G mutation (light purple squares). Symbols represent mean ± range from two independent experiments (error bars are smaller than the size of the symbols), and lines represent nonlinear fit of a one-site binding model to the experimental data.

**Fig. S6.**
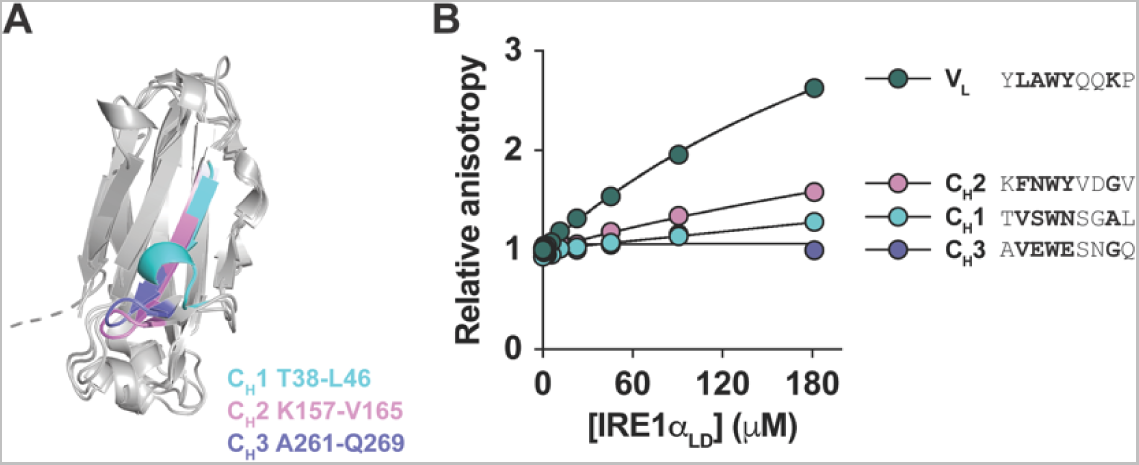
(A) Cartoon representation of human IgG heavy chain C_H_1, C_H_2, and C_H_3 domain superposition. Locations of peptides used for binding studies are colored on the structures. (B) Response curves for binding of IRE1α_LD_ to peptides derived from V_L_, C_H_1, C_H_2, and C_H_3 domains. Symbols represent measures from a single experiment, and lines represent nonlinear fit of a one-site binding model.

**Fig. S7.**
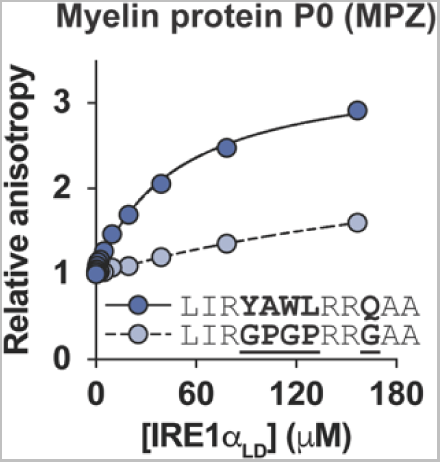
Response curves for binding of IRE1α_LD_ to WT (dark blue) and GPGPxxG mutant (light blue) peptides derived from myelin protein P0 (MPZ) measured by fluorescence anisotropy. Symbols represent mean ± SEM from three independent experiments, and lines represent nonlinear fit of one-site binding model.

**Fig. S8.**
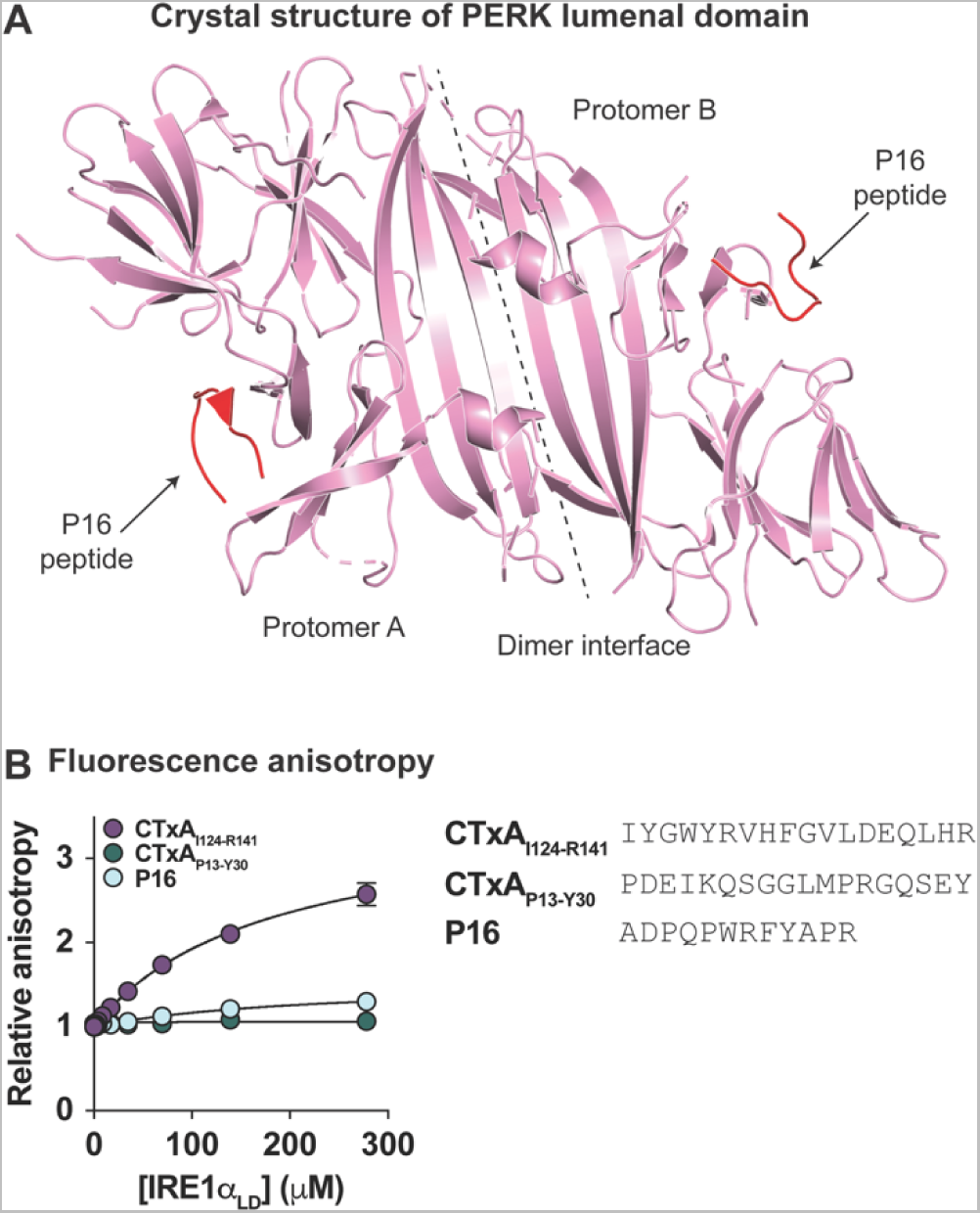
(A) Cartoon representation of crystal structure of PERK lumenal domain (pdb 5v1d, (*20*)). Location of the dimer interface and the exogenous P16 peptide (in red) are indicated. (B) Response curve for binding of IRE1α_LD_ to P16, CTxAI124-R141, and CTxAP13-Y30 peptides measured by fluorescence anisotropy. Symbols represent mean values from 2 independent experiments (CTxA peptides) or measures from a single experiment (P16 peptide).

**Table S1.**
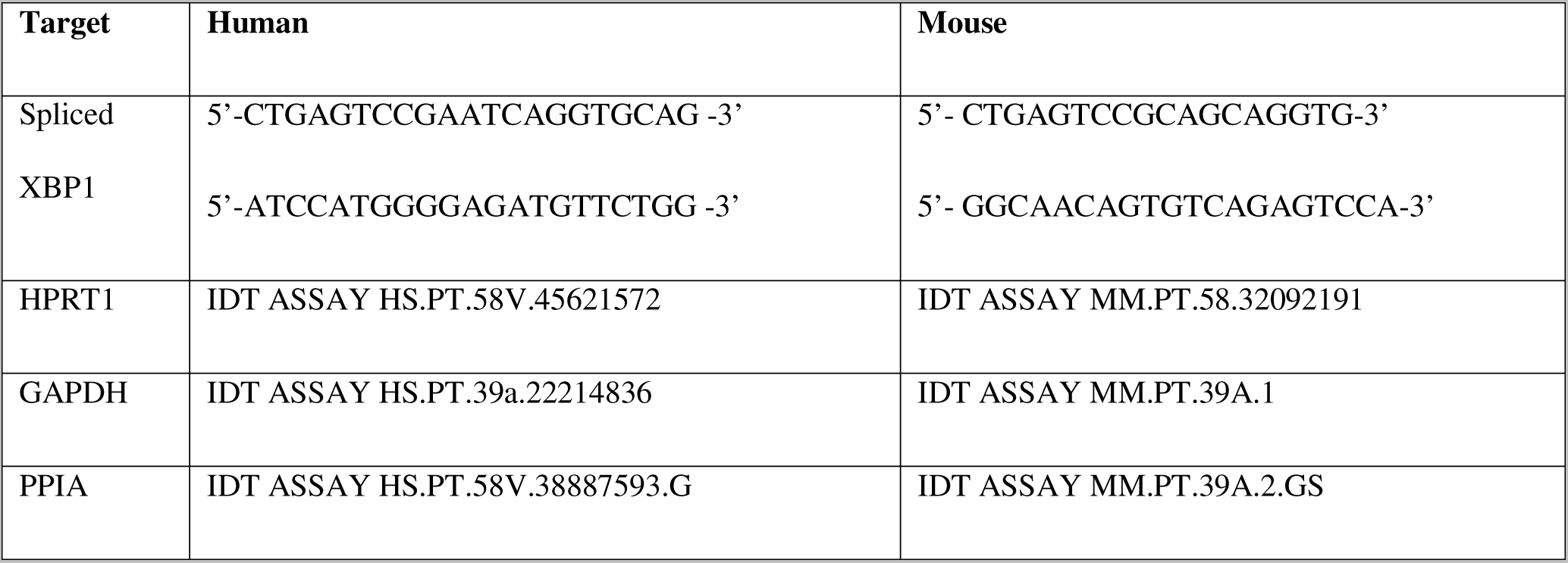
Primers used for expression analysis by qPCR

## References and Notes

1. W. Tirasophon, A. A. Welihinda, R. J. Kaufman, A stress response pathway from the endoplasmic reticulum to the nucleus requires a novel bifunctional protein kinase/endoribonuclease (Ire1p) in mammalian cells. Genes Dev 12, 1812–1824 (1998).

2. X. Z. Wang, H. P. Harding, Y. Zhang, E. M. Jolicoeur, M. Kuroda, D. Ron, Cloning of mammalian Ire1 reveals diversity in the ER stress responses. Embo j 17, 5708–5717 (1998).

3. G. E. Karagöz, D. Acosta-Alvear, P. Walter, The Unfolded Protein Response: Detecting and Responding to Fluctuations in the Protein-Folding Capacity of the Endoplasmic Reticulum. Cold Spring Harb Perspect Biol 11, (2019).

4. S. Preissler, D. Ron, Early Events in the Endoplasmic Reticulum Unfolded Protein Response. Cold Spring Harb Perspect Biol 11, (2019).

5. V. Belyy, I. Zuazo-Gaztelu, A. Alamban, A. Ashkenazi, P. Walter, Endoplasmic reticulum stress activates human IRE1α through reversible assembly of inactive dimers into small oligomers. Elife 11, (2022).

6. A. Bertolotti, Y. Zhang, L. M. Hendershot, H. P. Harding, D. Ron, Dynamic interaction of BiP and ER stress transducers in the unfolded-protein response. Nat Cell Biol 2, 326–332 (2000).

7. M. Calfon, H. Zeng, F. Urano, J. H. Till, S. R. Hubbard, H. P. Harding, S. G. Clark, D. Ron, IRE1 couples endoplasmic reticulum load to secretory capacity by processing the XBP-1 mRNA. Nature 415, 92–96 (2002).

8. H. Yoshida, T. Matsui, A. Yamamoto, T. Okada, K. Mori, XBP1 mRNA is induced by ATF6 and spliced by IRE1 in response to ER stress to produce a highly active transcription factor. Cell 107, 881–891 (2001).

9. K. Lee, W. Tirasophon, X. Shen, M. Michalak, R. Prywes, T. Okada, H. Yoshida, K. Mori, R. J. Kaufman, IRE1-mediated unconventional mRNA splicing and S2P-mediated ATF6 cleavage merge to regulate XBP1 in signaling the unfolded protein response. Genes Dev 16, 452–466 (2002).

10. J. Grootjans, A. Kaser, R. J. Kaufman, R. S. Blumberg, The unfolded protein response in immunity and inflammation. Nat Rev Immunol 16, 469–484 (2016).

11. C. Hetz, K. Zhang, R. J. Kaufman, Mechanisms, regulation and functions of the unfolded protein response. Nat Rev Mol Cell Biol 21, 421–438 (2020).

12. M. Wang, R. J. Kaufman, Protein misfolding in the endoplasmic reticulum as a conduit to human disease. Nature 529, 326–335 (2016).

13. N. Amin-Wetzel, L. Neidhardt, Y. Yan, M. P. Mayer, D. Ron, Unstructured regions in IRE1α specify BiP-mediated destabilisation of the luminal domain dimer and repression of the UPR. Elife 8, (2019).

14. G. E. Karagoz, D. Acosta-Alvear, H. T. Nguyen, C. P. Lee, F. Chu, P. Walter, An unfolded protein-induced conformational switch activates mammalian IRE1. Elife 6, (2017).

15. O. Guttman, A. Le Thomas, S. Marsters, D. A. Lawrence, L. Gutgesell, I. Zuazo-Gaztelu, J. M. Harnoss, S. M. Haag, A. Murthy, G. Strasser, Z. Modrusan, T. Wu, I. Mellman, A. Ashkenazi, Antigen-derived peptides engage the ER stress sensor IRE1α to curb dendritic cell cross-presentation. J Cell Biol 221, (2022).

16. A. Sundaram, S. Appathurai, R. Plumb, M. Mariappan, Dynamic changes in complexes of IRE1α, PERK, and ATF6α during endoplasmic reticulum stress. Mol Biol Cell 29, 1376–1388 (2018).

17. J. J. Credle, J. S. Finer-Moore, F. R. Papa, R. M. Stroud, P. Walter, On the mechanism of sensing unfolded protein in the endoplasmic reticulum. Proc Natl Acad Sci U S A 102, 18773–18784 (2005).

18. B. M. Gardner, P. Walter, Unfolded proteins are Ire1-activating ligands that directly induce the unfolded protein response. Science 333, 1891–1894 (2011).

19. A. V. Korennykh, P. F. Egea, A. A. Korostelev, J. Finer-Moore, C. Zhang, K. M. Shokat, R. M. Stroud, P. Walter, The unfolded protein response signals through high-order assembly of Ire1. Nature 457, 687–693 (2009).

20. P. Wang, J. Li, J. Tao, B. Sha, The luminal domain of the ER stress sensor protein PERK binds misfolded proteins and thereby triggers PERK oligomerization. J Biol Chem 293, 4110–4121 (2018).

21. N. L. Wernick, D. J. Chinnapen, J. A. Cho, W. I. Lencer, Cholera toxin: an intracellular journey into the cytosol by way of the endoplasmic reticulum. Toxins (Basel) 2, 310–325 (2010).

22. J. A. Cho, A. H. Lee, B. Platzer, B. C. S. Cross, B. M. Gardner, H. De Luca, P. Luong, H. P. Harding, L. H. Glimcher, P. Walter, E. Fiebiger, D. Ron, J. C. Kagan, W. I. Lencer, The unfolded protein response element IRE1α senses bacterial proteins invading the ER to activate RIG-I and innate immune signaling. Cell Host Microbe 13, 558–569 (2013).

23. Note: In follow-up studies after the publication of (29), we found evidence that the CTx-induced inflammatory response by IRE1α was not reproducible and as such that paper was retracted (Cell Host Microbe 218, 571 (2018)). The fundamental finding that CTx activated IRE1α, however, was always found to be true as documented here.

24. C. H. Tang, S. Chang, A. W. Paton, J. C. Paton, D. I. Gabrilovich, H. L. Ploegh, J. R. Del Valle, C. C. Hu, Phosphorylation of IRE1 at S729 regulates RIDD in B cells and antibody production after immunization. J Cell Biol 217, 1739–1755 (2018).

25. B. C. Cross, P. J. Bond, P. G. Sadowski, B. K. Jha, J. Zak, J. M. Goodman, R. H. Silverman, T. A. Neubert, I. R. Baxendale, D. Ron, H. P. Harding, The molecular basis for selective inhibition of unconventional mRNA splicing by an IRE1-binding small molecule. Proc Natl Acad Sci U S A 109, E869–878 (2012).

26. B. Tsai, C. Rodighiero, W. I. Lencer, T. A. Rapoport, Protein disulfide isomerase acts as a redox-dependent chaperone to unfold cholera toxin. Cell 104, 937–948 (2001).

27. V. Hung, N. D. Udeshi, S. S. Lam, K. H. Loh, K. J. Cox, K. Pedram, S. A. Carr, A. Y. Ting, Spatially resolved proteomic mapping in living cells with the engineered peroxidase APEX2. Nat Protoc 11, 456–475 (2016).

28. K. M. Bernardi, M. L. Forster, W. I. Lencer, B. Tsai, Derlin-1 facilitates the retro-translocation of cholera toxin. Mol Biol Cell 19, 877–884 (2008).

29. F. C. Nery, I. A. Armata, J. E. Farley, J. A. Cho, U. Yaqub, P. Chen, C. C. da Hora, Q. Wang, M. Tagaya, C. Klein, B. Tannous, K. A. Caldwell, G. A. Caldwell, W. I. Lencer, Y. Ye, X. O. Breakefield, TorsinA participates in endoplasmic reticulum-associated degradation. Nat Commun 2, 393 (2011).

30. J. M. Williams, T. Inoue, L. Banks, B. Tsai, The ERdj5-Sel1L complex facilitates cholera toxin retrotranslocation. Mol Biol Cell 24, 785–795 (2013).

31. A. Winkeler, D. Gödderz, V. Herzog, A. Schmitz, BiP-dependent export of cholera toxin from endoplasmic reticulum-derived microsomes. FEBS Lett 554, 439–442 (2003).

32. A. H. Pande, P. Scaglione, M. Taylor, K. N. Nemec, S. Tuthill, D. Moe, R. K. Holmes, S. A. Tatulian, K. Teter, Conformational instability of the cholera toxin A1 polypeptide. J Mol Biol 374, 1114–1128 (2007).

33. B. Goins, E. Freire, Thermal stability and intersubunit interactions of cholera toxin in solution and in association with its cell-surface receptor ganglioside GM1. Biochemistry 27, 2046–2052 (1988).

34. R. G. Zhang, D. L. Scott, M. L. Westbrook, S. Nance, B. D. Spangler, G. G. Shipley, E. M. Westbrook, The three-dimensional crystal structure of cholera toxin. J Mol Biol 251, 563–573 (1995).

35. J. Zhou, C. Y. Liu, S. H. Back, R. L. Clark, D. Peisach, Z. Xu, R. J. Kaufman, The crystal structure of human IRE1 luminal domain reveals a conserved dimerization interface required for activation of the unfolded protein response. Proc Natl Acad Sci U S A 103, 14343–14348 (2006).

36. Note: The construct used for crystallization of IRE1α_LD_ in (*35*)included up to residue 398. Residues 358-368 were resolved in the crystal structures, while the surrounding residues 307-357 and 369-398 included in the construct were not resolved in the structure.

37. X. Lu, C. Li, C. Li, P. Li, E. Fu, Y. Xie, F. Jin, Heat-Labile Enterotoxin-Induced PERK-CHOP Pathway Activation Causes Intestinal Epithelial Cell Apoptosis. Front Cell Infect Microbiol 7, 244 (2017).

38. L. Echavarría-Consuegra, G. M. Cook, I. Busnadiego, C. Lefèvre, S. Keep, K. Brown, N. Doyle, G. Dowgier, K. Franaszek, N. A. Moore, S. G. Siddell, E. Bickerton, B. G. Hale, A. E. Firth, I. Brierley, N. Irigoyen, Manipulation of the unfolded protein response: A pharmacological strategy against coronavirus infection. PLoS Pathog 17, e1009644 (2021).

39. F. Rashid, E. E. Dzakah, H. Wang, S. Tang, The ORF8 protein of SARS-CoV-2 induced endoplasmic reticulum stress and mediated immune evasion by antagonizing production of interferon beta. Virus Res 296, 198350 (2021).

40. J. Jung, E. Dudek, M. Michalak, The role of N-glycan in folding, trafficking and pathogenicity of myelin oligodendrocyte glycoprotein (MOG). Biochim Biophys Acta 1853, 2115–2121 (2015).

41. N. Amin-Wetzel, R. A. Saunders, M. J. Kamphuis, C. Rato, S. Preissler, H. P. Harding, D. Ron, A J-Protein Co-chaperone Recruits BiP to Monomerize IRE1 and Repress the Unfolded Protein Response. Cell 171, 1625–1637.e1613 (2017).

42. A. Bakunts, A. Orsi, M. Vitale, A. Cattaneo, F. Lari, L. Tadè, R. Sitia, A. Raimondi, A. Bachi, E. van Anken, Ratiometric sensing of BiP-client versus BiP levels by the unfolded protein response determines its signaling amplitude. Elife 6, (2017).

43. M. Vitale, A. Bakunts, A. Orsi, F. Lari, L. Tadè, A. Danieli, C. Rato, C. Valetti, R. Sitia, A. Raimondi, J. C. Christianson, E. van Anken, Inadequate BiP availability defines endoplasmic reticulum stress. Elife 8, (2019).

44. B. Weber, M. J. Brandl, M. D. Pulido Cendales, C. Berner, T. Pradhan, G. M. Feind, M. Zacharias, B. Reif, J. Buchner, A single residue switch reveals principles of antibody domain integrity. J Biol Chem 293, 17107–17118 (2018).

45. S. Mukherjee, S. P. Pondaven, K. Hand, J. Madine, C. P. Jaroniec, Effect of amino acid mutations on the conformational dynamics of amyloidogenic immunoglobulin light-chains: A combined NMR and in silico study. Sci Rep 7, 10339 (2017).

46. M. Karlsson, C. Zhang, L. Méar, W. Zhong, A. Digre, B. Katona, E. Sjöstedt, L. Butler, J. Odeberg, P. Dusart, F. Edfors, P. Oksvold, K. von Feilitzen, M. Zwahlen, M. Arif, O. Altay, X. Li, M. Ozcan, A. Mardinoglu, L. Fagerberg, J. Mulder, Y. Luo, F. Ponten, M. Uhlén, C. Lindskog, A single-cell type transcriptomics map of human tissues. Sci Adv 7, (2021).

47. Human Protein Atlas, proteinatlas.org.

48. A. M. Reimold, N. N. Iwakoshi, J. Manis, P. Vallabhajosyula, E. Szomolanyi-Tsuda, E. M. Gravallese, D. Friend, M. J. Grusby, F. Alt, L. H. Glimcher, Plasma cell differentiation requires the transcription factor XBP-1. Nature 412, 300–307 (2001).

49. N. N. Iwakoshi, A. H. Lee, P. Vallabhajosyula, K. L. Otipoby, K. Rajewsky, L. H. Glimcher, Plasma cell differentiation and the unfolded protein response intersect at the transcription factor XBP-1. Nat Immunol 4, 321–329 (2003).

50. A. L. Shaffer, M. Shapiro-Shelef, N. N. Iwakoshi, A. H. Lee, S. B. Qian, H. Zhao, X. Yu, L. Yang, B. K. Tan, A. Rosenwald, E. M. Hurt, E. Petroulakis, N. Sonenberg, J. W. Yewdell, K. Calame, L. H. Glimcher, L. M. Staudt, XBP1, downstream of Blimp-1, expands the secretory apparatus and other organelles, and increases protein synthesis in plasma cell differentiation. Immunity 21, 81–93 (2004).

51. E. Cloots, M. S. Simpson, C. De Nolf, W. I. Lencer, S. Janssens, M. J. Grey, Evolution and function of the epithelial cell-specific ER stress sensor IRE1β. Mucosal Immunol 14, 1235–1246 (2021).

52. M. F. Flajnik, M. Kasahara, Origin and evolution of the adaptive immune system: genetic events and selective pressures. Nat Rev Genet 11, 47–59 (2010).

53. S. Janssens, B. Pulendran, B. N. Lambrecht, Emerging functions of the unfolded protein response in immunity. Nat Immunol 15, 910–919 (2014).

54. D. T. Rutkowski, R. S. Hegde, Regulation of basal cellular physiology by the homeostatic unfolded protein response. J Cell Biol 189, 783–794 (2010).

55. W. I. Lencer, J. B. de Almeida, S. Moe, J. L. Stow, D. A. Ausiello, J. L. Madara, Entry of cholera toxin into polarized human intestinal epithelial cells. Identification of an early brefeldin A sensitive event required for A1-peptide generation. J Clin Invest 92, 2941–2951 (1993).

56. T. Iwawaki, R. Akai, Analysis of the XBP1 splicing mechanism using endoplasmic reticulum stress-indicators. Biochem Biophys Res Commun 350, 709–715 (2006).

57. M. J. Grey, E. Cloots, M. S. Simpson, N. LeDuc, Y. V. Serebrenik, H. De Luca, D. De Sutter, P. Luong, J. R. Thiagarajah, A. W. Paton, J. C. Paton, M. A. Seeliger, S. Eyckerman, S. Janssens, W. I. Lencer, IRE1β negatively regulates IRE1α signaling in response to endoplasmic reticulum stress. J Cell Biol 219, (2020).

58. M. J. Grey, H. De Luca, D. V. Ward, I. A. Kreulen, K. Bugda Gwilt, S. E. Foley, J. R. Thiagarajah, B. A. McCormick, J. R. Turner, W. I. Lencer, The epithelial-specific ER stress sensor ERN2/IRE1β enables host-microbiota crosstalk to affect colon goblet cell development. J Clin Invest 132, (2022).

59. H. Miyoshi, T. S. Stappenbeck, In vitro expansion and genetic modification of gastrointestinal stem cells in spheroid culture. Nat Protoc 8, 2471–2482 (2013).

60. J. Li, S. Han, H. Li, N. D. Udeshi, T. Svinkina, D. R. Mani, C. Xu, R. Guajardo, Q. Xie, T. Li, D. J. Luginbuhl, B. Wu, C. N. McLaughlin, A. Xie, P. Kaewsapsak, S. R. Quake, S. A. Carr, A. Y. Ting, L. Luo, Cell-Surface Proteomic Profiling in the Fly Brain Uncovers Wiring Regulators. Cell 180, 373–386.e315 (2020).

61. S. Qin, H. X. Zhou, meta-PPISP: a meta web server for protein-protein interaction site prediction. Bioinformatics 23, 3386–3387 (2007).

